# The allosteric landscape of the Src kinase

**DOI:** 10.1101/2024.04.26.591297

**Authors:** Antoni Beltran, Andre J. Faure, Ben Lehner

## Abstract

Enzymes catalyze the reactions of life and are the targets of most small molecule drugs. Most drugs target conserved enzyme active sites, often causing problems of specificity and toxicity. Targeting allosteric sites can increase specificity, overcome resistance mutations, and allow fine-tuning of activity. However, most enzymes have no known allosteric sites and methods do not exist to comprehensively identify them. Here we present a general and fast approach to chart allosteric communication in enzymes and apply it to the Src kinase to produce the first comprehensive map of negative and positive allosteric control of an enzymatic activity. Allostery in the Src kinase domain is pervasive, anisotropic, partially predictable, and modulated by regulatory domains. Multiple surface pockets of Src are allosterically active and so genetically-prioritized for the development of inhibitory and activating drugs. Using this approach it should be possible to chart global allosteric maps of many kinases and other enzymes important for medicine and biotechnology.

**Highlights:**

- First comprehensive map of negative and positive allosteric control of an enzymatic activity, the Src kinase.
- Allosteric communication is pervasive, distance dependent, and anisotropic.
- Allostery is conserved and modulated in the presence of the Src regulatory domains.
- Genetic prioritization of druggable surface pockets for Src inhibition and activation.
- Allosteric maps can now be constructed for many medically and industrially important kinases and enzymes.

## Introduction

Enzymes catalyze reactions important for nearly all biological processes from gene expression to metabolism, signaling, and neuronal computation^1^. Enzymes are also the largest class of targets of small molecule therapeutics^2^. Important examples include nucleotide metabolism and DNA replication enzymes in cancer and viral infections, lipid metabolism enzymes in metabolic disease, enzymes regulating synaptic transmission in neurological and psychiatric illnesses, and protein kinases in cancer and many other diseases^2,3^. Most drugs targeting enzymes have an orthosteric mechanism-of-action, inhibiting activity by binding directly to a protein’s active site. However, the active sites of enzymes are typically highly conserved, making it difficult to specifically target one enzyme without also inhibiting many others from the same family. This is the case for protein kinases that catalyze the transfer of phosphate groups from ATP to target proteins to regulate their activity. The human genome encodes 538 protein kinases, and orthosteric inhibitors targeting the ATP binding site typically inhibit tens to hundreds of different kinases^4–6^. The non-specific inhibition of off-target enzymes by orthosteric drugs is a major cause of drug toxicity^2^.

One strategy to increase drug specificity and reduce toxicity is to target allosteric sites^7^. A key mechanism of enzyme control is allosteric regulation whereby catalytic activity is regulated by small molecule, metabolite or macromolecule binding - or post translational modification - at a distant site^8^. Such long-range allosteric transmission of information in proteins underlies nearly all biological regulation and has been deemed ‘the second secret of life’^8^. Allosteric sites are less conserved between proteins and allosteric drugs can therefore have higher specificity and reduced toxicity than orthosteric drugs^7^. Targeting allosteric sites can also result in activation or in quantitative modulation of activity, for example tuning the response to an agonist^9^.

Most enzymes and most proteins, however, have no known allosteric sites to target therapeutically. Indeed, the prevalence of allostery across the protein universe is unknown. A key reason for this is the lack of methods that systematically quantify allosteric communication^10–12^. Fast and general methods to globally quantify allosteric regulation of enzyme activity would transform our ability to understand, predict, target, and engineer allosteric control.

Due to their pivotal role in many physiological and pathological processes, protein kinases now constitute the second most important class of small molecule drug targets^13,14^. In cancer, driver mutations hyperactivate kinases and kinase signaling and effective kinase inhibitors have been developed^13,14^, including inhibitors of BCR-ABL in chronic myeloid leukemia^15^, BRAF and MEK inhibitors in melanoma^16^, EGFR and ALK inhibitors in lung cancer^17^ and EGFR/HER2 inhibitors in breast cancer^18^. Most kinase inhibitors developed to date have an orthosteric mechanism-of-action, targeting the highly conserved ATP binding pocket, resulting in limited specificity^13,14^. Moreover, the efficacy of many kinase inhibitors in cancer is short-lived due to selection for resistance mutations^13,14,19^.

To increase specificity and overcome resistance mutations, allosteric inhibitors have been developed against several kinases^20^. These include GNF-2, a highly selective inhibitor targeting the myristate-binding site of BCR-ABL^21^, trametinib and selutinib, MEK1/2 inhibitors that bind a pocket located in the vicinity of the ATP binding site and helix αC^22^, and MK-2206, an AKT inhibitor that binds at the interface between its PH and kinase domains^23^, among many others^20^. However, although allostery is likely part of the regulation of all kinases^24^, most kinases have no known allosteric sites^25^. Rather, every kinase has tens of different surface pockets that could be potentially targeted with small molecules and it is not known which of these pockets - if any - is allosterically active^26^. In short, we do not know which pockets in which kinases are allosterically active and which should be prioritized for drug development.

Here, we present a general approach that can be used to globally quantify allosteric regulation of enzyme activities. We use the method to produce the first comprehensive map of positive and negative allosteric control of an enzymatic activity, the Src kinase. Src was the first discovered oncogene^27^, and is a significant drug target in cancer due to its role in regulating cell growth, cell migration, and angiogenesis^28^. Using *in vivo* assays, we quantify the effects of more than 50,000 single and double mutant variants on the activity and abundance of the Src kinase for the kinase domain in isolation and in the context of the full-length enzyme, totalling 30 separate pooled selection assays and more than 200,000 variant effect measurements. We use thermodynamic modeling to disambiguate the effects of mutations on kinase activity and abundance, thereby revealing complete allosteric maps of the Src kinase domain alone and in the context of its regulatory domains. Allosteric control of Src is pervasive, distance dependent, anisotropic, and reasonably well predicted by sequence and structural features. These maps enable the comprehensive analysis of the positive and negative allosteric control of the druggable surface of an enzyme, allowing unbiased genetic prioritization of many novel Src surface pockets for drug discovery. The approach that we describe will allow comprehensive allosteric maps to be charted for many other kinases and therapeutically and biotechnologically important enzymes.

## Results

### A general strategy to map allosteric regulation of enzyme activity

Most enzymes have no known allosteric sites to target for drug discovery or biotechnological control. We therefore conceived a general strategy that uses genetics to comprehensively quantify allosteric regulation of enzyme activity.

Our approach has three steps: first, the effects of mutations on catalytic activity are quantified; second, the effects of the same mutations on the abundance of the enzyme are measured; and third, changes in activity that cannot be accounted for by changes in concentration are quantified by fitting a model to the data. Due to the non-linear relationships between the energetic effects of mutations and molecular phenotypes, we quantify the effects of mutations in multiple protein variants with different abundance and/or activities to constrain model fitting, allowing the underlying causal effects of mutations to be determined^12,29–31^. Both enzymatic activity^32–35^ and protein abundance^36–40^ can be quantified using many different selection assays. Thus, although here we demonstrate the approach using a selection assay suitable for one particular class of enzymes (protein kinases) and one particular method to quantify protein abundance, this general approach can be used to quantify allosteric regulation in any enzyme, provided that both activity and abundance can be quantified at scale.

### Measuring activity and abundance of Src protein kinase variants at scale

To demonstrate the feasibility of the approach we apply it here to the human oncoprotein Src. Src was the first discovered tyrosine kinase, and the first identified, cloned, and sequenced oncogene^27,41^. The enzymatic activity of variants of Src is easy to measure using a highly-validated cellular toxicity assay where the inhibition of cellular growth is directly proportional to the amount of Src-induced protein phosphorylation^42–44^. The solubility of Src can also be quantified in the same cells using a highly-validated protein abundance selection assay, abundancePCA (aPCA), that uses protein fragment complementation to quantify soluble protein concentration over at least three orders of magnitude^12,30,40^.

To generate sufficient data for model fitting, we constructed five overlapping libraries of Src variants that together cover all possible single mutants in the kinase domain (KD), with each variant present in at least 10 different genetic backgrounds. The genetic backgrounds were selected to provide a range of different Src activities due to either abundance or catalytic activity changes (see Methods). In total, the five libraries contain a total of 54,455 genotypes. We quantified the kinase activity and abundance of each genotype in 30 separate pooled selection assays. First, we quantified kinase activity in triplicate using kinase-dependent impairment of cell growth^42–45^ (Figure 1a,b). Activity scores were highly reproducible for all five library tiles (median Pearson’s *r* = 0.90) and strongly correlated with Src-dependent tyrosine phosphorylation^42,43^ (*r* = −0.89, Figure 1c, Figure S1a,b). Second, we quantified the cellular abundance of Src kinase domain variants using aPCA^30,40^ (Figure 1d,e). Abundance measurements were also reproducible (median *r* = 0.75, Figure S1a,c) and correlated with the in vivo abundance of SRC variants measured by western blotting^42,43^ (*r* = 0.66, Figure 1f). Measured fitness scores for all genotypes and their associated errors are provided in Supplementary Table 1.

**Figure 1:**
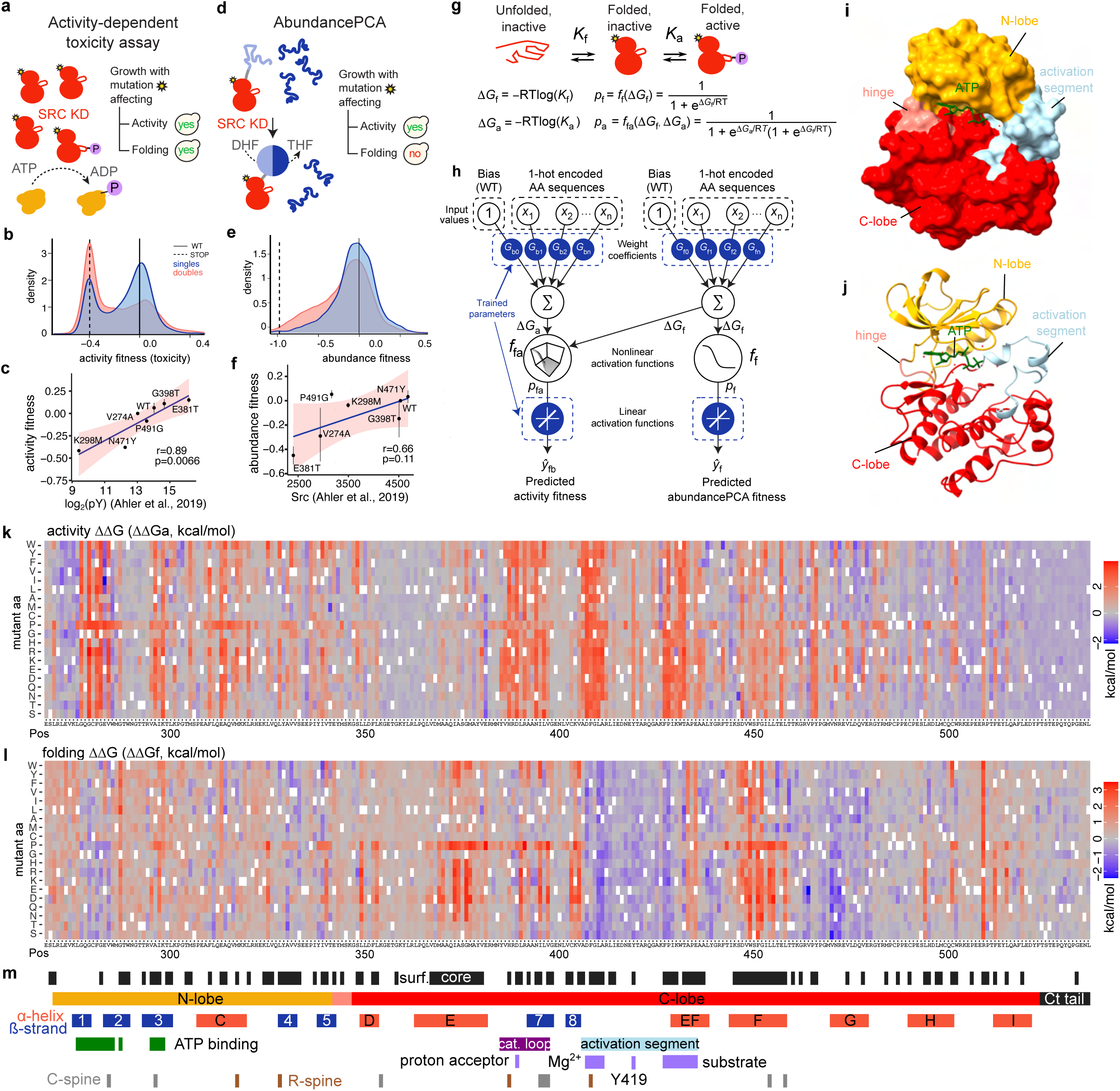
Mapping the energetic and allosteric landscapes of enzymes. **a.** Overview of the toxicity selection assay to measure the protein kinase activity of Src kinase domain variants at scale. yes, yeast growth; no, yeast growth defect. **b.** Activity fitness distribution of single and double mutants of Src. **c.** Correlation of activity fitness measurements to in vivo phosphotyrosine levels^43^. **d.** Overview of the abundancePCA (aPCA) selection assay to measure in vivo abundance of Src kinase domain variants at scale. yes, yeast growth; no, yeast growth defect. DHF, dihydrofolate; THF,tetrahydrofolate. **e.** aPCA fitness distribution of single and double mutants of Src. **f.** Correlation of aPCA fitness measurements to in vivo Src abundance^43^. **g**. Three-state equilibrium and corresponding thermodynamic model. ΔG_f_, Gibbs free energy of folding; ΔG_a_, Gibbs free energy of the active state; Kf, folding equilibrium constant; Ka, inactive-active state equilibrium constant; pf, fraction folded; pfa, fraction folded and active; ff, nonlinear function of ΔG_f_; ffa, nonlinear function of ΔG_f_ and ΔG_a_; R, gas constant; T, temperature in Kelvin. **h**. Neural network architecture used to fit thermodynamic models to the toxicity and abundance data (bottom, target and output data), thereby inferring the causal changes in free energy of folding and binding associated with single amino acid substitutions (top, input values). **i, j**. 3D structure of the Src kinase domain (Protein Data Bank (PDB) ID: 2SRC). **k, l**. Heat maps showing inferred changes in activity free energies (k, ΔΔG_a_) and folding free energies (l, ΔΔG_f_). **m**, Sequence and annotation of Src. Locations of individual secondary structure elements and functional regions were obtained from ^48,71^.

### Quantifying changes in activity not caused by changes in abundance

To quantify the changes in Src activity that are not accounted for by changes in the cellular abundance of Src, we fit a simple phenomenological ‘enzyme folding and activation’ model to the data (see Methods). In this model, the folding of Src is explicitly modeled using a two-state thermodynamic model in which the protein can exist in two states - unfolded (U) and folded (F), with the Gibbs free energy of folding, ΔG_f_, determining the partitioning of Src molecules between the two states according to the Boltzmann distribution^12,30,31^ (Figure 1g-m). All other biophysical changes that alter Src kinase activity are quantified using a second pseudo free energy which we refer to as the activity energy, ΔG_a_ (Figure 1g-m). The model is formally equivalent to a 3-state model with unfolded (U), folded inactive (F), and folded active (A) states. However, the active state modeled here is phenomenological, and designed to quantify all changes in activity not accounted for by changes in total soluble protein abundance. Although shifts in the equilibrium between inactive and active kinase conformations will be captured as changes in ΔG_a_, so too will other molecular mechanisms that affect catalytic activity (kcat) and substrate affinity (Km) independently of the conformational state of Src.

Fitting the enzyme folding and activation model to the data provides very good prediction of the abundance and activity of double mutants (median percent variance explained 82.6% for activity and 68.8% for abundance, evaluated by 10-fold cross validation, Figure S1d) and a marked improvement over a 2-state (unfolded, folded active) model (Figure S1e). In contrast, increasing the complexity of the model to four states did not improve performance (Figure S1e). In total, our dataset quantifies folding and activity energies for all of the 5,111 possible single substitution variants in the Src KD (Supplementary Table 2).

### Stability of the kinase fold

Our data provides the first comprehensive measurements of how mutations affect the *in vivo* abundance of the protein kinase fold and one of the largest sets of abundance measurements for any protein (Figure 2a, Supplementary Movie 1). The Src KD is composed of two structurally and functionally distinct subdomains - the N- and C-lobes - with the active site located in the cleft between the two. The N-lobe is mostly composed of beta strands and contains the ATP binding site, whereas the C-lobe is mostly alpha helical and ends in a disordered C-terminal tail that regulates the conformational state of the kinase^46–48^. Mutations have a wide range of effects, with many destabilizing (703 strongly destabilizing mutations with ΔΔG_f_>0.5, p<0.05, z-test) and a large number of moderately stabilizing variants (468 with ΔΔG_f_<0, p<0.05, z-test).

**Figure 2:**
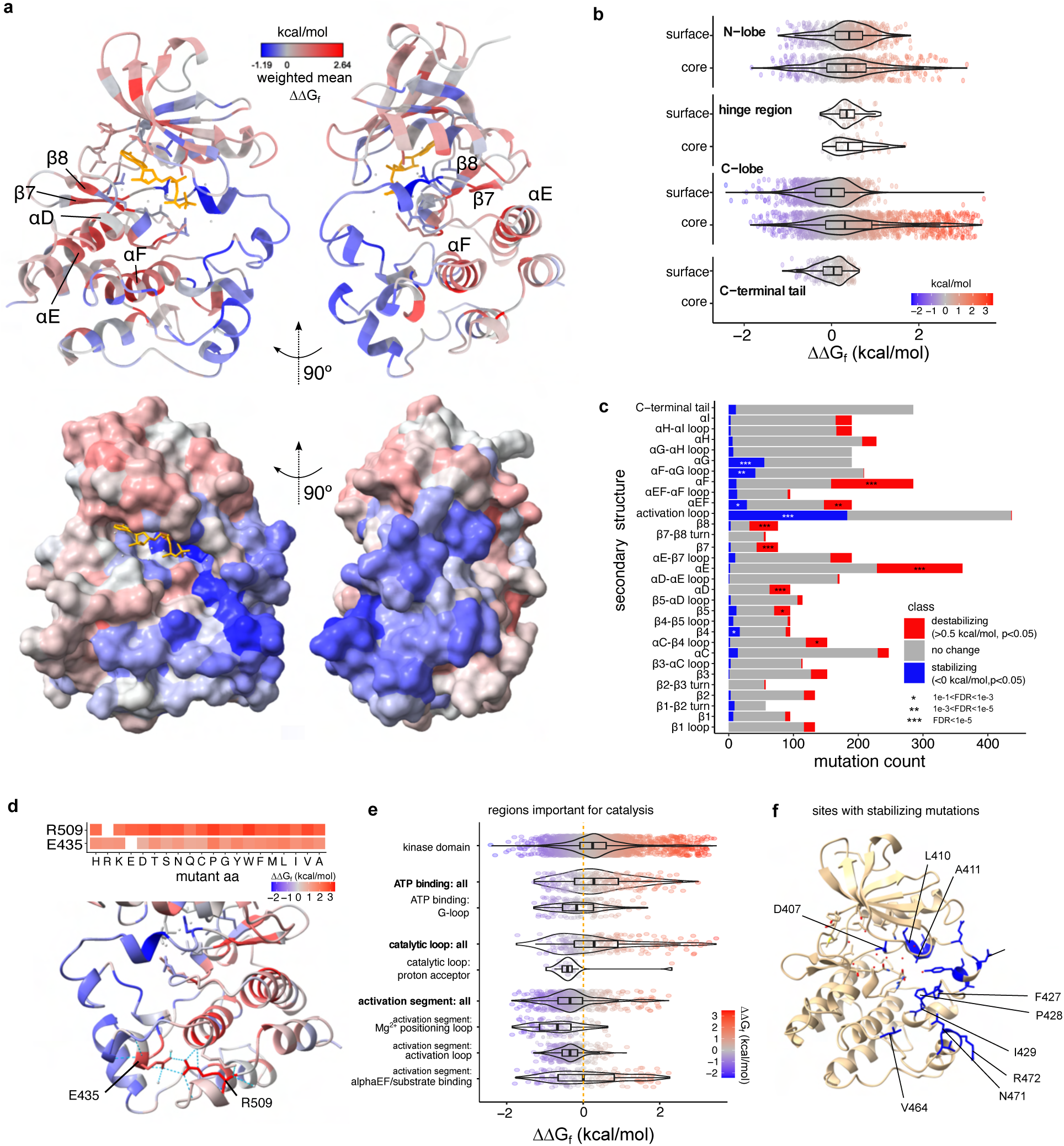
The folding landscape of the Src kinase domain. **a.** Structure of the Src KD coloured by the per-site weighted mean ΔΔG_f_ (PDB ID: 2SRC). The secondary structure elements most enriched in destabilizing mutations are annotated. **b**. ΔΔG_f_ distributions across the core and surface of the two kinase lobes. **c**. Enrichment of destabilizing and stabilizing mutations in secondary structure elements of Src (Fisher’s exact test). **d**. ΔΔG_f_ of mutations in the E435-R509 salt bridge. **e**. ΔΔG_f_ distributions in catalytically important functional regions of Src. **f**. Sites enriched in mutations increasing in vivo stability of the Src KD (FDR<0.1).

Destabilizing mutations across the KD show strong structural biases. Mutations in the core of the KD (relative solvent accessible surface area, rSASA<0.25) are much more likely to be destabilizing than mutations on the surface (odds ratio (OR) = 10.10, p = 8.02e-116, Fisher’s exact test (FET), Figure 2a,b). This enrichment is much stronger for the C-lobe (OR = 25.81, p = 3.10e-110, FET) than for the N-lobe (OR = 2.30, p = 3.78e-6, FET). Mutations in the C-terminal tail only have mild effects on stability (Figure 2b). In addition, individual secondary structure elements are particularly sensitive to mutation, with the hydrophobic αF helix buried in the C-lobe core most critical for stability (Figure 2c). The αE helix that contacts αF extensively, αEF, and β7 and β8 that are packed against αE are also highly enriched in destabilizing mutations (Figure 2a,c). Together these elements form the main stabilizing core of the kinase fold (Figure 2a,c). Mutations to proline are most likely to be destabilizing, especially in helices and beta strands (Figure 1l, Figure S2a)^49^. Finally, of all long-range side-chain to side-chain structural contacts, the salt bridge connecting R509 in helix αI with E435 in αEF is the most sensitive to mutation (maximum mean ΔΔG_f_ of residue pairs), with no substitutions tolerated in either residue (Figure 2d). Mutations in E435-R509 are 1.91-fold more destabilizing than mutations in the next most sensitive contact (the K298-E313 salt bridge), 3.33-fold more than the average of all salt bridges, and 5.45-fold more than the average of all side-chain to side-chain contacts.

The two structurally distinct lobes of the Src kinase domain thus also contribute differentially to the *in vivo* stability of the domain, with the more dynamic ATP-binding N-lobe displaying a higher tolerance to mutagenesis than the larger and more compact C-lobe.

### Stabilizing mutations and molecular frustration

Quantifying mutational effects across a range of destabilized genetic backgrounds allows us to identify a total of 468 mutations that moderately increase the cellular abundance of the Src KD, which we define as *in vivo* stabilizing variants (ΔΔG_f_<0, p<0.05, z-test, Figure 2c). These variants are distributed in 118 positions in the Src KD. Examination of the spatial distribution of stabilizing mutations in the structure reveals a striking continuous surface of sites spanning the ATP binding site, the activation loop and the αG helix (Figure 2a,c). Stabilizing variants are enriched in residues essential for catalysis (Figure 2e), including the G-loop that positions ATP for catalysis (Figure 2e, OR = 1.45, p = 0.15, FET), the proton acceptor D389 in the catalytic loop (OR = 2.66, p = 0.09, FET), and the activation segment (OR = 9.34,p = 5.72e-84, FET). The activation segment contains over half of all stabilizing mutations, particularly in the Mg2+ positioning loop (OR = 14.62,p = 2.01e-32, FET), with L410 and A411 individually enriched (FDR<0.1, FET, Figure 2f), and in the substrate positioning loop (OR = 5.04,p = 3.21e-17, FET) (Figure 2e,f), with many stabilizing mutations in F427, P428 and I429 (FDR<0.1, FET, Figure 2f). In addition, V464 in the αF-αG loop, and N471 and R472 in αG are particularly enriched (FDR<0.1, FET, Figure 2e,f).

Many residues in the kinase domain of Src are thus highly frustrated, with side chains important for catalytic activity compromising the *in vivo* stability of the protein^50,51^. This may reflect both the need for catalytically important side chains that are not optimal for fold stability and solubility, and the requirement for flexibility for substrate binding and product release during catalysis.

### The Src active site

Our data provides the first complete map of the effects of mutations on protein kinase activity independently of their effects on protein abundance (Figure 3a, Supplementary Movie 2). In total, we identified 1273 mutations that modulate kinase activity more than can be accounted for by changes in Src abundance (|ΔΔG_a_|>0.5, FDR<0.1, z-test). 1227 of these mutations are inhibitory and 46 are activating. Mutations in the active site – residues that directly contact ATP, Mg2+ or the substrate peptide phosphosite – are overwhelmingly detrimental to kinase activity, with 225/247 decreasing activity (ΔΔG_a_>0.5, OR = 39.39, p = 1.63e-118, FET, Figure 3b,c). Inactivating variants are strongly enriched in the ATP binding site (OR = 3.78, p = 1.41e-24, FET), the catalytic loop (HRD motif, OR = 5.85, p = 1.32e-39, FET), the Mg2+ positioning loop (DFG motif, OR = 32.43, p = 5.92e-44, FET) and the substrate positioning loop (OR = 9.37, p = 1.00e-42, FET) (Figure 3b). Mutations in the beta sheet that forms the top surface of the ATP binding pocket have a striking alternating pattern of mutational effects, with substitutions of side chains pointing towards the nucleotide detrimental for activity (Figure 3d).

**Figure 3:**
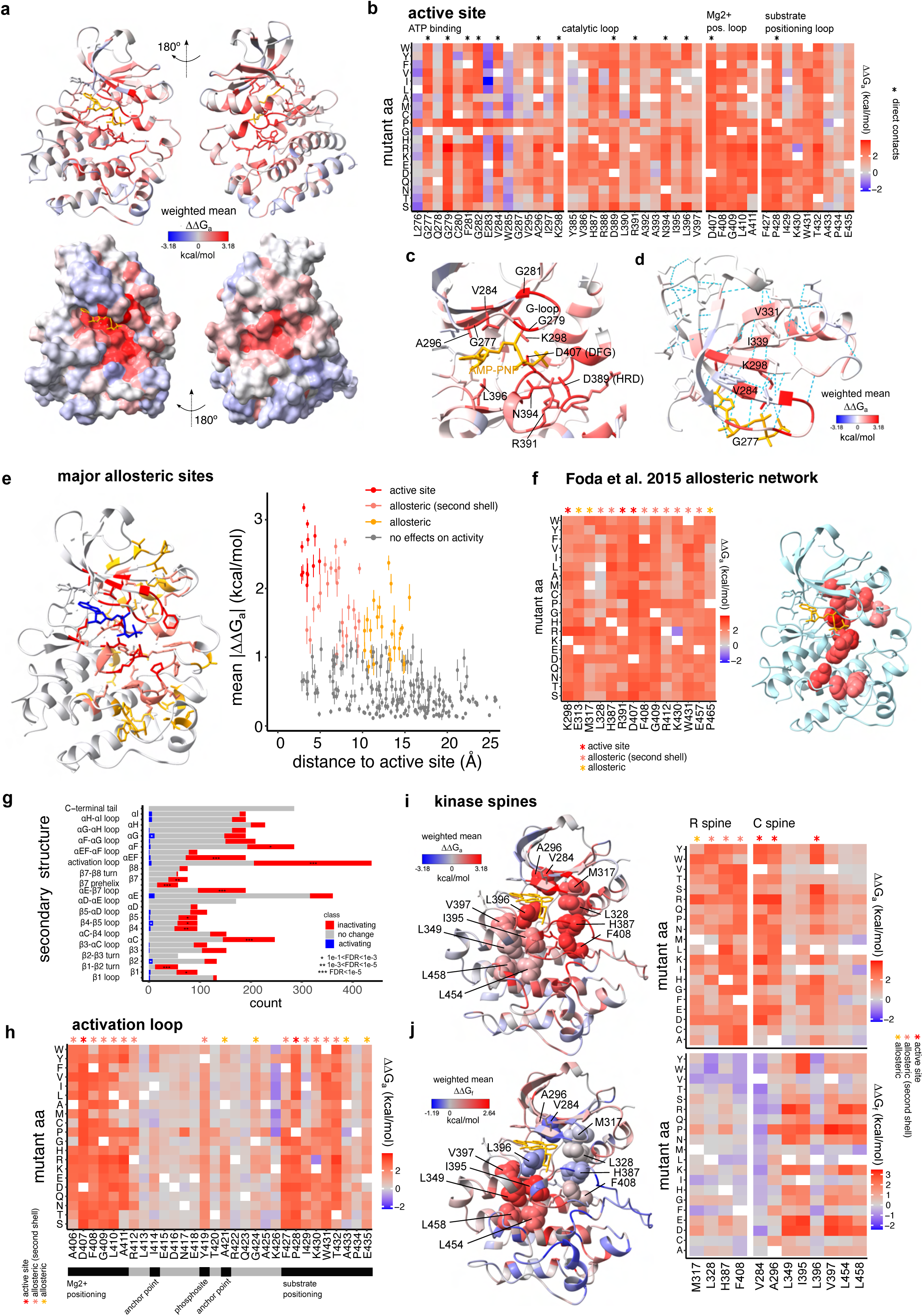
The allosteric landscape of the Src kinase domain. **a.** Structure of the Src KD coloured by the per-site weighted mean ΔΔG_a_ (PDB ID: 2SRC). **b**. Heat map showing ΔΔG_a_ of residues in the Src regions forming the active site. Residues directly contacting the nucleotide, Mg^2+^ or the substrate peptide are labeled with stars. **c**. The active site of Src (PDB ID: 2SRC). **d**. Beta sheet forming the top surface of the ATP binding pocket (N-lobe). **e**. Relationship between per-site averaged ΔΔG_a_ and the minimum heavy atom distance to the active site. Error bars represent the standard error of the mean. **f**. Heat map showing ΔΔG_a_ of mutations in residues that are part of a previously described allosteric network connecting ATP and substrate binding sites^52^. **g**. Enrichment of inactivating and activating mutations in secondary structure elements (Fisher’s exact test). **h,i**. Src structures and heatmaps showing ΔΔG_a_ (d) and ΔΔG_f_ (e) of mutations in the catalytic (C) and regulatory (R) spines of Src.

### Major allosteric sites

We identified 1048 mutations in 174 sites located outside of the active site that modulate kinase activity (|ΔΔGa|>0.5,FDR<0.1, z-test). 1002 of these allosteric mutations are inhibitory and 46 activate the kinase. We define major allosteric sites as residues outside the active site that are enriched for these mutations (Figure 3e,f). By this definition, Src has 47 major allosteric sites (OR>2, FDR<0.1, FET): 13 in the N-lobe and 34 in the C-lobe (Supplementary Movie 3).

Strikingly, 23/47 major allosteric sites are second shell sites directly contacting residues in the active site. The major allosteric sites also include all 11 non-active site residues previously predicted to be part of an allosteric network that communicates between the substrate and ATP binding sites of Src (Figure 3f). This network was predicted via analysis of changes in electrostatic and hydrophobic contacts between active and inactive conformations in molecular dynamics simulations^52^. Of these 11 previously predicted allosteric positions, 8 are second shell residues. The predicted allosteric network is very strongly enriched for mutations with large ΔΔG_a_ (OR = 20.75, p = 3.77e-80, FET, excluding active site residues, and OR = 10.53, p = 1.13e-16, FET, excluding active site and second shell residues), with all individual residues enriched at least 6-fold for allosteric mutations (Figure 3f).

### Inhibitory allosteric mutations

Defining major inhibitory allosteric sites as sites enriched for inhibitory mutations (OR>2, FDR<0.1, FET) identifies 47 positions, all of which are also major allosteric sites. Inhibitory allosteric mutations are enriched in several structural elements (Figure 3g), including helix αC, consistent with its conformational change upon Src activation^46,48^. Inhibitory mutations are concentrated in residues on the inner surface of helix αC, including E313 that engages in a salt bridge with K298 in the active state (Figure 3f), the R-spine (see below) residue M317, and the hydrophobic residues F310, A314 and L320. The β4 and β5 strands located between αC and the active site are also enriched for allosteric mutations, but to a lesser extent (Figure 3g). Inhibitory allosteric mutations are also abundant in the activation loop (Figure 3g,h) including in Y419, which locks Src in the active state when phosphorylated^48,53^ (Figure 3h); in αEF, that functions in substrate positioning; and in αG, in hydrophobic residues that contact αEF.

Finally, inhibitory allosteric mutations are enriched in the αF helix that acts as an anchor for the catalytic (C) and regulatory (R) ‘spines’ (Figure 3h,i). The C- and R-spines are two groups of residues that are not contiguous in the primary sequence of kinases but form a bipartite hydrophobic core in catalytically active kinases^54,55^. Mutations in all R-spine residues have strong inactivating effects independently of their effects on abundance (Figure 3h). In contrast, only the C-spine residues in direct contact with ATP (A296, V284, L396) are enriched in inactivating mutations (Figure 3h). Mutations in the rest of the C-spine sites have small effects on ΔG_a_, and their strong effects on kinase activity at the fitness level are almost fully explained by a loss of fold stability (Figure 3i).

### Activating allosteric mutations

In total, 11 residues outside of the active site are enriched for activating mutations, which we define as major activating allosteric sites (OR>1, p<0.05, FET, Figure S3). In the N-lobe, major activating allosteric sites include the gatekeeper residue T341^56^, as well as its neighboring residue Y343. E283 that flanks the G-loop and forms a salt bridge with K275 to constrain the conformation of the G-loop, and E335 in the β4-β5 loop are also major activating allosteric sites. In the C-lobe, major activating allosteric sites are located in the αF pocket (E381, T511 and Y514) and in surface-exposed residues of helix αG (V470, D476, E479) (Figure S3). Finally, activating mutations are also enriched in K426, which is adjacent to residues that position the substrate peptide Y for phosphorylation. Interestingly, all substitutions of K426 to hydrophobic residues increase activity, with the exception of K426Y which is inhibitory (Figure 3h, Figure S3).

### The distance dependence of allosteric regulation

We next considered the spatial organization of allosteric mutations. The average effect of mutations on Src activity is much stronger closer to the Src active site (Figure 4a, Figure S4a). Indeed, considering all 252 residues in the Src KD, there is an exponential decay of mutational effects on activity away from the active site (Figure 4a) with a decay rate k = −0.093 ± 0.005 Å^-1^ (95% confidence interval), corresponding to a 50% reduction of allosteric effects over a distance d_1/2_ = 7.45Å (Figure 4a). The distance dependence is, on average, similar in the N- and C-lobes of the KD, despite their structural differences (k = −0.092 ± 0.011 Å^-1^ for the N-lobe and k = −0.110 ± 0.006 Å^-1^ for the C-lobe) (Figure 4b). Interestingly, in contrast to what is observed for inactivating mutations (ΔΔG_a_>0), whose effects scale with distance (k = −0.083 ± 0.006 Å^-1^) (Figure 4d), the distance dependence of activating mutations is extremely weak (k = −0.017 ± 0.007 Å^-1^) (Figure 4c). This effect is not driven by the smaller effect size of activating mutations as inactivating mutations with matched effect sizes have a decay almost twice as fast (median k = −0.030 Å^-1^, simulation p = 1.3e-3, n=10,000 subsamples, Figure S5c-e). This suggests different mechanisms underlie inhibitory and activating mutations.

**Figure 4:**
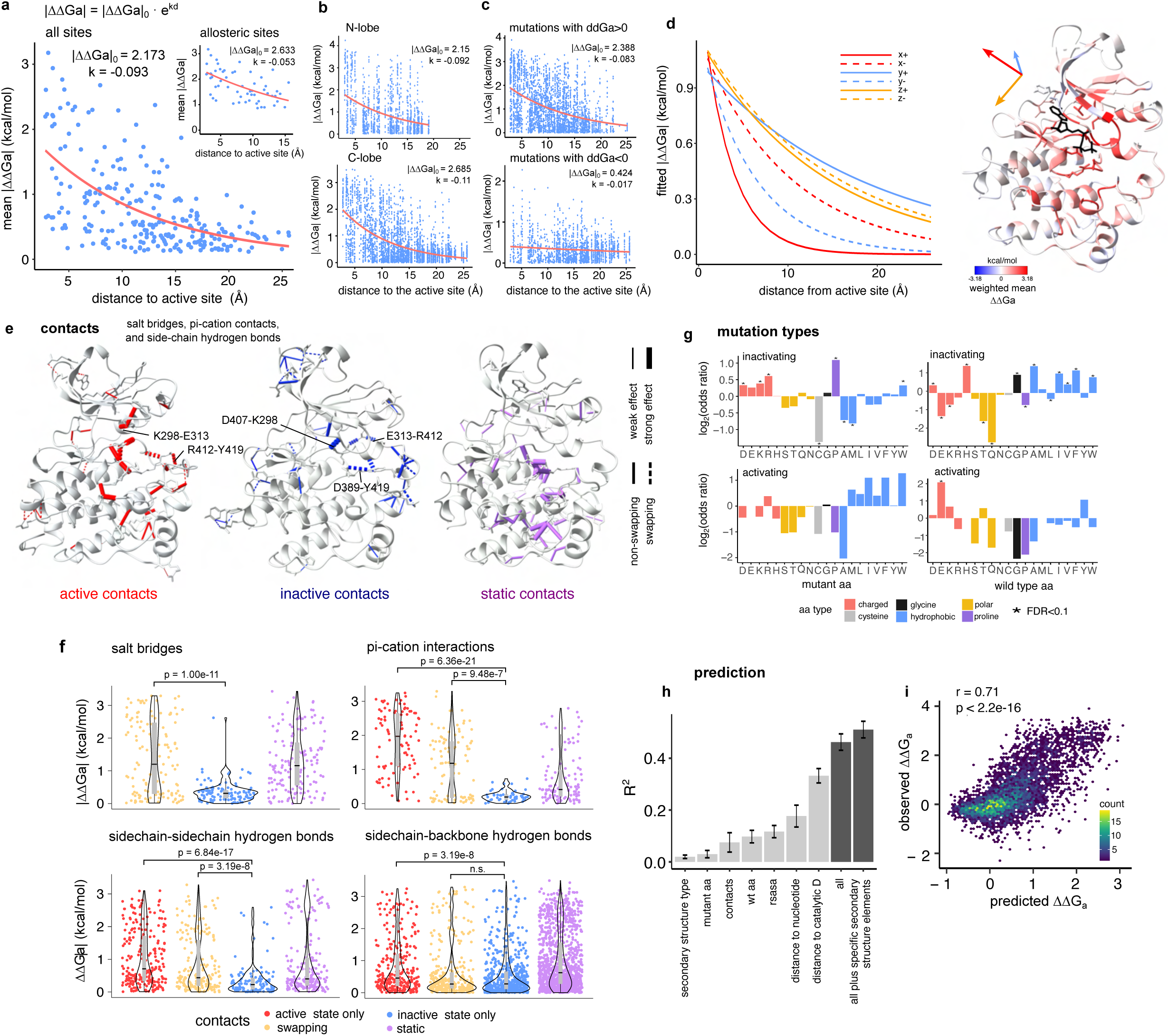
Allosteric communication is anisotropic and enriched in dynamic structural contacts. **a.** Exponential decay fits to the relationship between per-site averaged |ΔΔG_a_| to the minimum heavy atom distance to the active site, for all sites (main) and for active and allosteric sites (inset). |ΔΔG_a_|_0_=starting ΔΔG_a_ at distance=0 (active site), k=decay rate, d=distance from the active site. **b**. Exponential decay fits of |ΔΔG_a_| with distance to the active site for the N- and C-lobes. **c**. Exponential decay fits of |ΔΔG_a_| with distance to the active site for inactivating (ΔΔG_a_>0) and activating (ΔΔG_a_<0) mutations. **d**. Exponential decay fits of |ΔΔG_a_| with distance to the active site in the three orthogonal spatial directions, positive and negative. Decay was calculated in each spatial direction for residues located at a distance less than 10Å from the active site in the two remaining directions. **e**. Src structures depicting non-covalent contacts specific to the active state of Src (PDB ID: 1Y57), specific to the inactive state of Src (PDB ID: 2SRC), and present in both. Salt bridges, side chain to side chain and side chain to backbone hydrogen bonds, and pi-cation contacts are shown. The thickness of the contact is proportional to the averaged ΔΔG_a_ of mutations in the two contacting residues. Contacts occurring in residues that swap contacting partners between active and inactive conformations (‘swapping residues’) are depicted as dashed lines. **f**. Distributions of ΔΔG_a_ of mutations in residues classified according to their contacting patterns in the active and inactive states of Src, for four contact types. Adjusted p-values of Wilcoxon rank sum tests to compare mutation effect distributions are shown. **g**. Enrichments of mutations in inactivating and activating groups according to the WT (mutations ‘from’) and mutated (mutations ‘to’) amino acid identities. **h**. R^2^ of single predictor linear models (light gray), and the final models incorporating all features (dark gray). R^2^ values were calculated on held out data in a 10-fold cross-validation strategy. Error bars are the standard deviations of the R^2^ values of the 10 folds. **i**. Correlation between observed and predicted ΔΔG_a_ for the model incorporating all features.

However, allosteric communication in the kinase domain is not isotropic, as mutations in particular residues and in particular directions are more likely to be allosteric at a given distance. To illustrate this, the distance dependence when only considering the major allosteric sites is much weaker (k = −0.053 ± 0.008 Å^-1^, d_1/2_ = 12.38Å) (Figure 4a, inset, Figure S4b). Major allosteric sites are not arranged uniformly throughout the kinase domain. Instead, they are spatially clustered and have higher connectivity than expected by chance (Figure S4f-i). Indeed, quantification of the allosteric decay rate from the active site in 3 orthogonal spatial axes (x,y,z, axes as defined in PDB ID: 2SRC) in the positive and negative directions (+,-) (Figure 4d, Figure S4j), reveals more effective transmission in the direction towards helix αC (y+, k = −0.053 ± 0.013 Å^-1^, d_1/2_ = 13.08 Å), and in the vertical axis of the KD (z+, k = −0.073 ± 0.009 Å^-1^, d_1/2_ = 9.50 Å and z-, k = −0.068 ± 0.013 Å^-1^, d_1/2_ = 10.19 Å), and less effective transmission towards the regulatory domain interaction surfaces (x+, k = - 0.325 ± 0.070 Å^-1^, d_1/2_ = 2.25 Å). Allosteric transmission also differs across secondary structure types, with faster decay for mutations occurring in beta strands, where mutations have smaller effects than expected given their distance to the active site (Figure S4k,l).

Inhibitory allosteric communication in the Src KD is thus strongly distance dependent, but also anisotropic: transmission efficiency is dependent on the direction of propagation, with at least a 6-fold difference in decay rates between the most and least efficient directions.

### Allosteric communication via dynamic non-covalent contacts

The conformation of the Src KD differs between its active and inactive states, with changes in the positioning of helix αC and the activation loop and multiple residue contact rearrangements in the active site and throughout the kinase domain^48,52,57^ (Figure 4e). Based on their contacts in active (PDB ID: 1Y57) and inactive (PDB ID: 2SRC) state structures of Src, we define four types of residues for different types of contacts (e.g. salt bridges, pi-cation interactions): active-only (engaging in contacts only in the active state), inactive-only (only in the inactive state), swapping (residues that have different contacts in the two states), and static (residues with the same contacts in both states) (Figure 4e).

Overall, mutations in active-only and swapping residues are more likely to affect the activity of the Src KD than those in inactive state-only residues (Figure 4f) .The differences in |ΔΔG_a_| are strongest when considering contacts between side chains, including salt bridges, pi-cation interactions, and side-chain to side-chain hydrogen bonds (Figure 4f). Swapping residues include those forming Src’s ‘electrostatic switch network’ of contacts that change during activation: D407-K298, E313-R412 and D389-Y419 in the inactive state, that break and rearrange into E313-K298 and R412-Y419 (Figure 4f). Mutations in these residues are extremely detrimental for Src activity (Figure 4f, Figure 1k). The allosteric map thus shows that residues with contacts that change upon activation are particularly important for Src activation and enables the prioritization of which of these dynamic contacts are most important for activation.

### Amino acid changes and allostery

We next considered different types of mutation across the domain. Mutations at histidine, glycine, and several hydrophobic residues (alanine, isoleucine, valine, phenylalanine, and tryptophan) are more likely to have inhibitory effects (FDR<0.1, FET) and substitutions to proline are the most likely to be inhibitory (FDR<0.1, FET, Figure 4g), as also observed for allosteric regulation of protein-protein interactions^12,30^. In Src, substitutions to charged residues (glutamate, lysine and arginine) and to tryptophan, the largest aa, are also more likely to be allosteric (Figure 4g). Substitutions to cysteine, alanine and methionine are the least likely to be allosteric (FDR<0.1, FET, Figure 4g). The smaller number of activating allosteric mutations makes analyses of their properties less powered. However, mutations at glutamate residues (FDR<0.1, FET) and substitutions to hydrophobic aa are more likely to be allosteric activating mutations (p = 7.6e-3, FET for hydrophobics as a group) (Figure 4g).

### Predicting allosteric mutations

We next asked how much of the variance in allostery (ΔΔG_a_) for mutations outside of the active site can be accounted for by sequence and structural features. We used linear modeling to predict ΔΔG_a_ from simple features: the minimum heavy atom distance of the mutated residue to the nucleotide (AMP-PNP in PDB structure 2SRC) and to the catalytic residue D389, the identity of the wild-type and mutant aa, solvent accessibility, contact type and dynamics (active-only, inactive-only, swapping, and static), and secondary structure element type. Distance to the catalytic site and to the nucleotide are the most predictive features when tested individually (Figure 4h). A linear model combining all predictors explains 46% of the variance in ΔΔGa (tested on held out data, 10-fold cross-validation, Figure 4h), which increases further to 51% when incorporating specific secondary structure elements as a feature (tested on held out data, 10-fold cross-validation, Figure 4h,i). Mutation effects on activity are thus reasonably well-predicted from simple structural features alone, illustrating the potential to predict allostery from sequence.

### Genetic prioritization of allosteric surface pockets

Most kinase inhibitors developed to-date target the highly conserved orthosteric ATP binding site, resulting in limited specificity^13,14^. Allosteric inhibitors have been successfully developed to increase specificity and overcome drug resistance mutations for several kinases including BCR-ABL^21^, BRAF, MEK^22^ and AKT^23^. However, each kinase has many potentially druggable surface pockets and, beyond a very small number of examples, it is not known which of these pockets are allosterically active in each kinase.

Structural analysis of the surface of Src (see Methods) identifies 28 unique small molecule fragment-binding hotspots (henceforth referred to as surface pockets) present in at least one of 15 different Src structures^25^ (Figure 5a, Figure S5). To prioritize these pockets for drug development, we used our comprehensive atlas to quantitatively rank these pockets based on their allosteric activity (Figure 5b).

**Figure 5:**
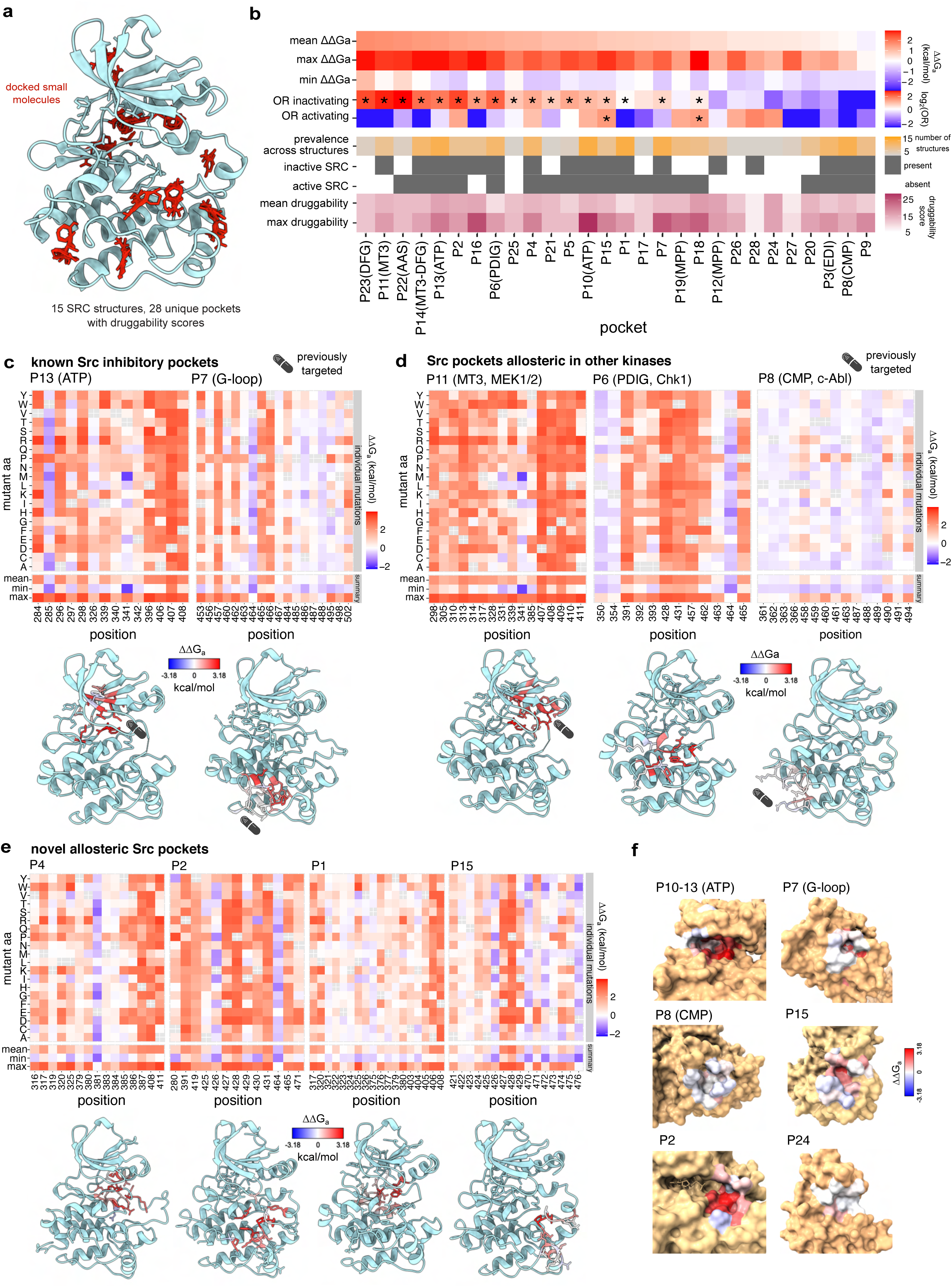
The druggable allosteric surface of Src. **a.** Overview of the Kinase Atlas pocket dataset for the Src kinase, consisting of 15 Src structures with multiple small molecule docking sites and their associated druggability scores. **b**. Summary of the regulatory impact and druggability properties of Src surface pockets. Mean ΔΔG_a_ = average of per-site averaged ΔΔG_a_ of all residues in the pocket. Max ΔΔG_a_ = maximum of per-site averaged ΔΔG_a_ of all residues in the pocket. Min ΔΔG_a_ = minimum of per-site averaged ΔΔG_a_ of all residues in the pocket. Pockets significantly enriched or depleted in activating and inactivating mutations (Fisher’s exact test FDR<0.05) are labeled with stars. **c,d,e**. Heatmaps showing ΔΔG_a_ of mutations in Src previously targeted pockets (c), in Src pockets homologous to pockets known to be allosteric and/or targeted by drugs in other kinases (d), and in novel Src pockets (e). The residues forming each pocket are highlighted in the structure of Src (PDB ID: 2SRC), coloured by their weighted mean ΔΔG_a_. Pockets previously targeted by drugs are labeled with a capsule pill. **f**. Surface view of representative examples of Src pockets, including highly druggable and allosterically active pockets (P10-13(ATP),P7, P15, P2), and highly druggable but allosterically inactive pockets (P24, P8(CMP)).

In total, 17 Src pockets are enriched for inhibitory mutations, with varying levels of enrichment (Fisher’s exact test, FDR<0.05, see Figure 5b for a rank). These inhibitory pockets include the orthosteric ATP binding site targeted by competitive inhibitors. Beyond the orthosteric site, two other surface pockets of Src have been targeted by small molecule inhibitors: the DFG pocket^58^, and P7^59^. Consistent with this, both of these pockets are enriched for allosteric inhibitory mutations in the allosteric map, genetically validating their regulatory potential (Figure 5c).

### New allosteric surface pockets

Across 538 protein kinases, 12 different pockets have been reported as potentially allosteric^25^. Some of these pockets are the binding sites of small molecule inhibitors, whereas others are physiological binding sites for other proteins and lipids, or dimerization interfaces^25^. Of these 12 pockets, 7 have a structurally homologous pocket in Src^25^ (Supplementary Table 8), but it is unknown how many of these – if any – are also allosterically active in Src.

In total, four of these 7 pockets are enriched for inhibitory allosteric mutations (FDR<0.05, FET, Figure 5b,d). Pockets that are allosteric in other kinases and also strongly allosteric in Src include P11, homologous to the MT3 pocket in MEK1/2 targeted by type III allosteric inhibitors^22^, P22, homologous to the AAS site in Aurora A, where one KD activates another through binding of its activation segment to this site^60^, and P6, homologous to the PDIG pocket in CHK1 close to the substrate binding site that is bound by small molecule inhibitors^61^ (Figure 5b,d). In contrast, three other pockets homologous to allosteric pockets in other kinases show little evidence of allostery in Src. These include pocket P3 (Figure 5b), homologous to the EDI site that is part of the EGFR dimerization interface and P8 (Figure 5b,d), homologous to the Bcr-Abl myristoyl pocket (CMP) that has been targeted by small molecule inhibitors^21^. Interestingly, three of the residues that form the myristoyl pocket in Abl are substituted by bulky residues in Src^62^, likely rendering the pocket unable to bind myristate^62^ and consistent with the pocket having little evidence of allosteric activity in our dataset.

Finally, we considered all the surface pockets of Src that, to our knowledge, have not been previously reported as allosteric in any kinases^25^. A total of 10 of these 17 pockets are enriched for allosteric mutations in Src (Figure 5b,e). All 10 of these novel pockets are enriched for inhibitory allosteric mutations and 2 of them (P15 and P18) are also enriched for activating allosteric mutations. These novel allosteric pockets include P4, P16 and P25 located between the allosteric αC helix and the active site, P1 formed by residues in the αC-β4 loop, αE and β8, P18 and P21 in the surface of the N-lobe beta sheet, and P2, P15 and P5 located on both sides of αEF and the substrate positioning loop (see Figure 5e for examples). Computational methods^63–66^ do not successfully predict the allosteric pockets in Src (Figure S7). A full list of all pockets with druggability and inhibitory and activatory allosteric scores is available in Supplementary table 3, and full surface views of the kinase domain depicting per-residue weighted mean and maximum weighted ΔΔG_a_ are available as Supplementary Movies 4 and 5, respectively.

The comprehensive mutational effect data therefore serves to genetically prioritize which of the many potentially druggable surface pockets of Src should be the focus for inhibitory and activatory drug discovery. The allosterically active pockets in Src include highly druggable novel pockets not previously demonstrated as allosteric in any kinases (Figure 5e).

### Modulation of the allosteric landscape by the Src regulatory domains

Src, like most kinases and eukaryotic proteins, is a multi-domain protein. In addition to the catalytic KD, Src contains two additional globular domains, SH2 and SH3, disordered linkers and the dynamic SH4 region. The non-catalytic domains of Src physically interact with the KD in its inactive conformation and inhibit activity^48,57,67^. To ask how the regulatory domains of Src affect allosteric communication in the catalytic domain, we repeated our abundance and activity selections for the same 54,455 Src variants in the context of the full length protein (Figure 6a,b). The selections were highly reproducible (median r = 0.93 for both assays, Figure S6a) and the fitness scores correlated well with independent in vivo activity (r = 0.89) and abundance (r = 0.80) measurements (Figure S6b,c).

**Figure 6:**
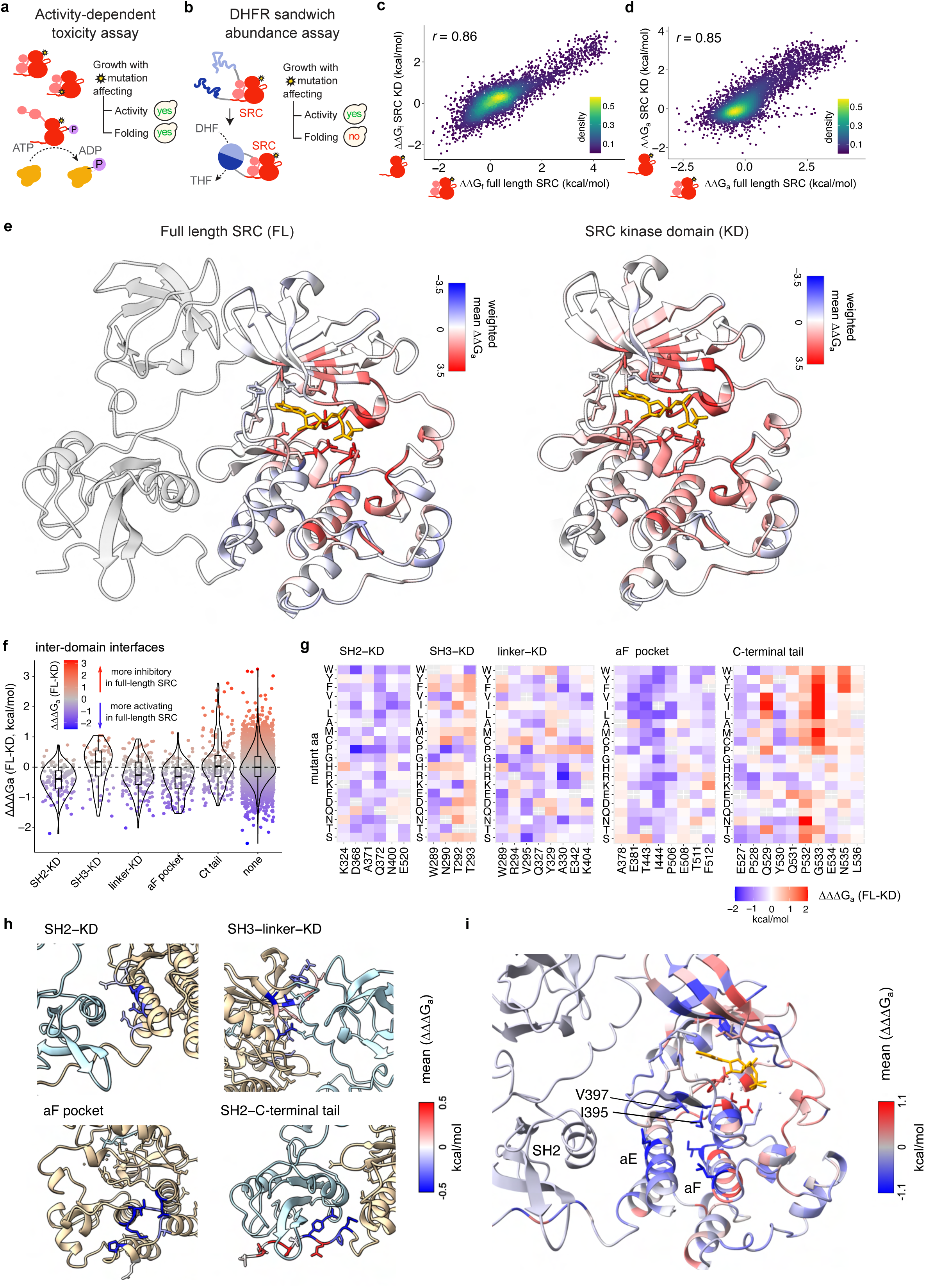
Impact of Src regulatory domains on the allosteric landscape. **a.** Overview of the toxicity selection assay to measure the protein kinase activity of full-length Src variants at scale. yes, yeast growth; no, yeast growth defect. **b.** Overview of the sandwich abundancePCA (sPCA) selection assay to measure in vivo abundance of full-length Src variants at scale. yes, yeast growth; no, yeast growth defect. DHF, dihydrofolate; THF,tetrahydrofolate. **c**. Correlation between inferred ΔΔG_f_ in full-length Src and ΔΔG_f_ in Src kinase domain. **d**. Correlation between inferred ΔΔG_a_ in full-length Src and ΔΔG_a_ in Src kinase domain. **e.** Structural view of the allosteric landscape of the Src kinase domain in the context of full length SRC (left structure) and in isolation (right panel). **f**. Distribution of changes in ΔΔG_a_ between full-length and kinase domain Src (ΔΔΔG_a_) at regulatory domain interfaces. **g**. Heatmaps showing changes in ΔΔG_a_ between full-length and kinase domain Src (ΔΔΔG_a_) at regulatory domain interfaces with the kinase domain. **h**. Src structures (PDB ID: 2SRC) with residues in regulatory domain interfaces coloured by mean ΔΔΔG_a_. **i**. Src structure coloured by mean ΔΔΔG, with the secondary structure elements and containing the C-lobe cluster of residues more activating in full-length Src labeled.

Fitting the same 3-state folding and activation model to the full length Src data allows us to quantitatively compare the effects of all 5,111 single amino acid substitutions on stability and activation in the presence and absence of the Src regulatory domains (Figure S6d-g, Supplementary Movie 6,7). Overall the mutational effects on folding energies and active state energies correlate very well between the kinase domain and full length Src constructs (*r* = 0.86 and *r* = 0.85, respectively, Figure 6c-e). The allosteric landscape of the kinase domain is therefore highly conserved in the presence of the regulatory domains (Figure 6e).

However, there are differences between the two allosteric landscapes. Activating mutations are more frequent in full-length Src than in the kinase domain alone (Figure 6e, Figure S6h), consistent with the regulatory domains having an overall inhibitory function^48,57,67^. In particular, mutations in the inter-domain surfaces with the SH2 domain and the SH2-KD linker have stronger activating effects in the full-length kinase (Figure 6f-h), consistent with these intra-molecular interactions inhibiting kinase activity. Mutations in the αF helix pocket, proposed to bind the SH4 region for additional inhibition of Src activity^43^, also more strongly activate full-length Src. These differences are not driven by changes in the effects of mutations on abundance between full-length Src and the KD alone, as fitting the 3-state folding and activation models exchanging the underlying abundance data does not affect the conclusions (Figure S6i).

Mutations in the dynamic C-terminal tail of Src also differ in their effects between full-length Src and the KD alone (Figure 6f-h). Mutations in Y530, the inhibitory phosphosite directly involved in the interaction with the SH2 domain, have stronger activating effects in full-length Src (ΔΔΔG_a_<-1, FDR<0.1, see Methods), consistent with a release of the inhibitory interaction. The adjacent E527, P528, and Q531 are similarly enriched for mutations with stronger activating effects in full-length Src. In contrast, Q529, P532, G533, and N535 are enriched for mutations with stronger inhibitory effects in full length Src (ΔΔΔG_a_>1, FDR<0.1) (Figure 6f-h). Inhibitory mutations in the C-terminal tail’s interface with the SH2 domain include many changes to hydrophobic and aromatic residues, which may act by increasing the affinity of the tail for the Src SH2 domain^48,57,67^.

Finally, spatial clustering of mutations with stronger activating effects in full-length Src (ΔΔΔG_a_< −1, FDR<0.1) reveals an additional cluster of residues in the C-lobe (Figure S6j, 6i). This cluster includes the SH2-KD interface on the outer surface of αE, along with the internal surface of αE, a second internal layer of residues in αF pointing towards αE, and the C-spine residues I395 and V397 (Figure 6i). Mutations in these sites at a distance from the SH2 domain interface are thus more activating in full length Src, potentially allosterically relieving inhibition by the SH2 domain.

## Discussion

We have presented here a general strategy to chart allosteric maps of enzymatic activities and have used a specific implementation to produce the first comprehensive map of negative and positive allosteric control of an enzyme, the Src kinase.

The Src kinase allosteric map provides a number of fundamental insights into kinase regulation and allostery.

First, allosteric communication occurs throughout the kinase domain.

Second, inhibitory allosteric effects are larger close to the active site and exponentially decay away from the active site with a half distance (d_1/2_) of 8.4 Å. This exponential decay is consistent with previous estimates of percolation and dissipation of protein structural perturbations^68^ and with evolutionary conservation gradients toward catalytic residues^69^.

Third, allosteric communication in Src is anisotropic, with 6-fold lower decay rates in some directions, and mutations in particular structural features such as αC and the activation loop particularly likely to have strong allosteric effects.

Fourth, residues with structural contacts that change upon kinase activation are particularly likely to be allosteric.

Fifth, particular types of mutations are more likely to be allosteric, including mutations to proline and to charged amino acids.

Sixth, mutations can both allosterically inhibit and activate Src. Activating allosteric mutations are both less abundant than inhibitory allosteric mutations and have a different spatial arrangement in the protein, without strong enrichment close to the active site, suggesting a different molecular mechanism.

Seventh, Src has many potentially druggable surface pockets but only a subset of these are strongly enriched for allosteric mutations. Our data quantitatively ranks 17 pockets to target to inhibit the enzyme and two to target to activate it. 10 of these pockets have not, to our knowledge, been previously identified as allosteric in any kinase. Existing computational methods^63–66^ do not successfully predict the allosteric sites and pockets in Src (Figure S7).

Eighth, allosteric communication in the Src kinase domain is highly conserved in the presence of the Src regulatory domains but a subset of mutations have stronger allosteric effects in the context of the regulatory domains, including substitutions at sites that contact the regulatory domains and potentially communicate their inhibition.

Ninth, allostery is, at least to a reasonable extent, predictable from sequence and structure.

We believe that the method presented here can be applied to many different enzymes to chart their regulatory allosteric maps. This adds to our previously developed method to chart allosteric regulation of protein-protein interactions^12,30^, and extends allosteric mapping to many additional classes of proteins important for medicine and biotechnology. The main requirement is for a selection assay that links genotype to enzymatic activity. Suitable selections exist for many different enzymatic activities, including microfluidic^33^ and growth-based complementation^35^ and toxicity assays^34^. Indeed, the specific assay employed in this study can likely be used to chart allosteric maps for many different protein kinases^44,70^, including tens of different human cancer genes. This will allow fundamental questions about the conservation and evolution of allosteric sites in protein families to be addressed. From a pharmaceutical perspective, these datasets will enable the systematic identification of novel surface pockets that are allosterically active and structurally accessible in only a subset of homologous proteins and thus contribute to the development of increasingly specific kinase and enzyme inhibitors. How the effects of mutations relate to the consequences of small molecule binding will be an interesting question to address in future studies.

We believe that mapping allosteric communication in selectable enzymes is an efficient experimental strategy that can be used to build allosteric datasets of sufficient size and diversity to both better understand the mechanistic principles of allosteric communication and to train machine learning models to predict allosteric sites. Indeed, we have shown here that simple structural and sequence features perform reasonably well for predicting allosteric mutations in Src. By quantifying allosteric communication in structurally diverse proteins, we should be able to train computational models to predict, therapeutically target, and engineer allosteric sites in any protein.

## Supplementary Figure legends

**Supplementary Figure 1:**
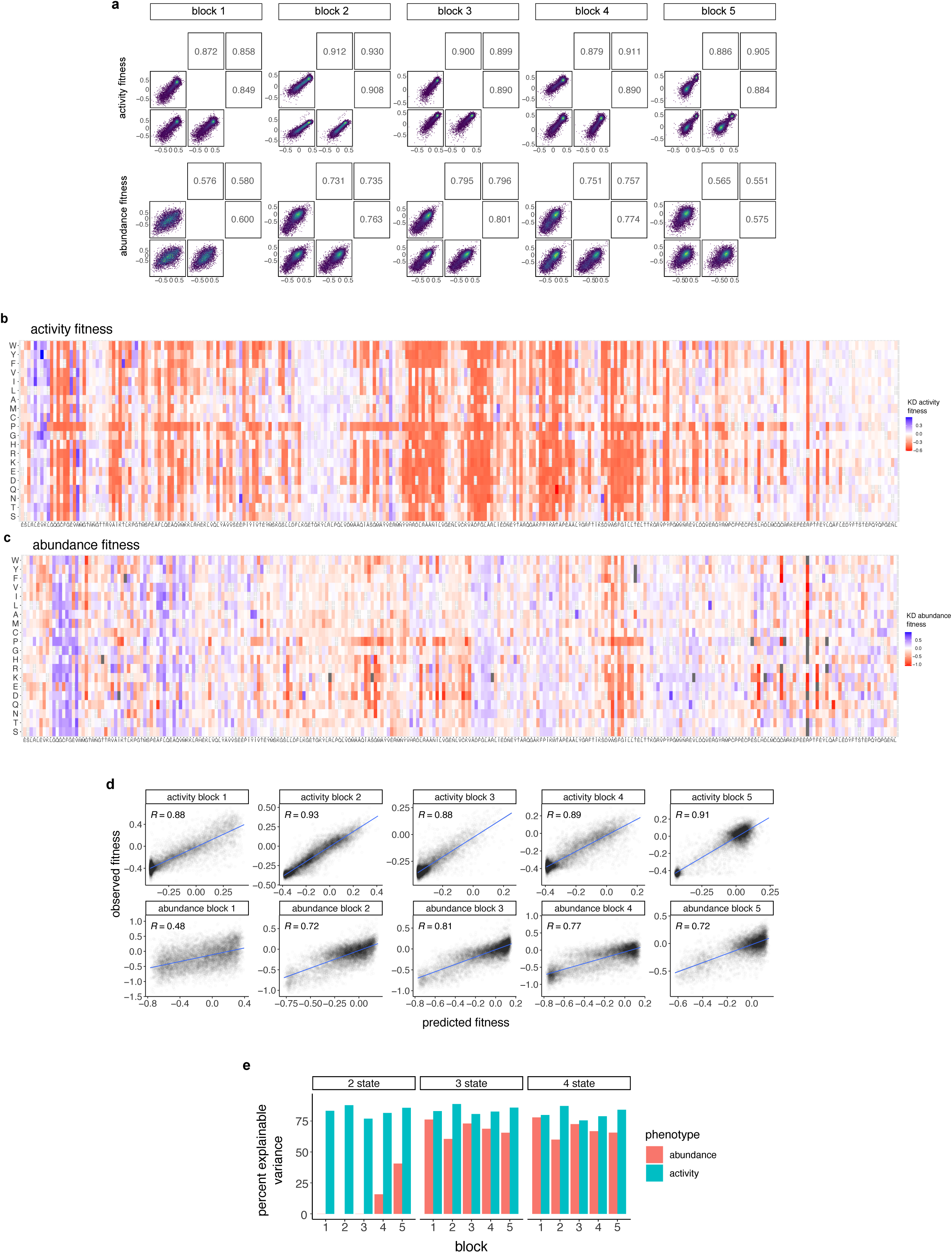
Selection assay reproducibility and thermodynamic model fitting evaluation. **a.** Fitness score replicate correlations for each of the 5 blocks of the Src kinase domain library, for the activity-dependent toxicity assay (top row) and abundancePCA (bottom row). **b-c.** Fitness score heatmaps for Src activity (b) and abundance (c). **d**. Correlations between observed fitness values and MoCHI fitness predictions for test set variants held out in any of the 10 training folds. **e**. Percentage of explainable variance captured by 2-state, 3-state, and 4-state models.

**Supplementary Figure 2:**
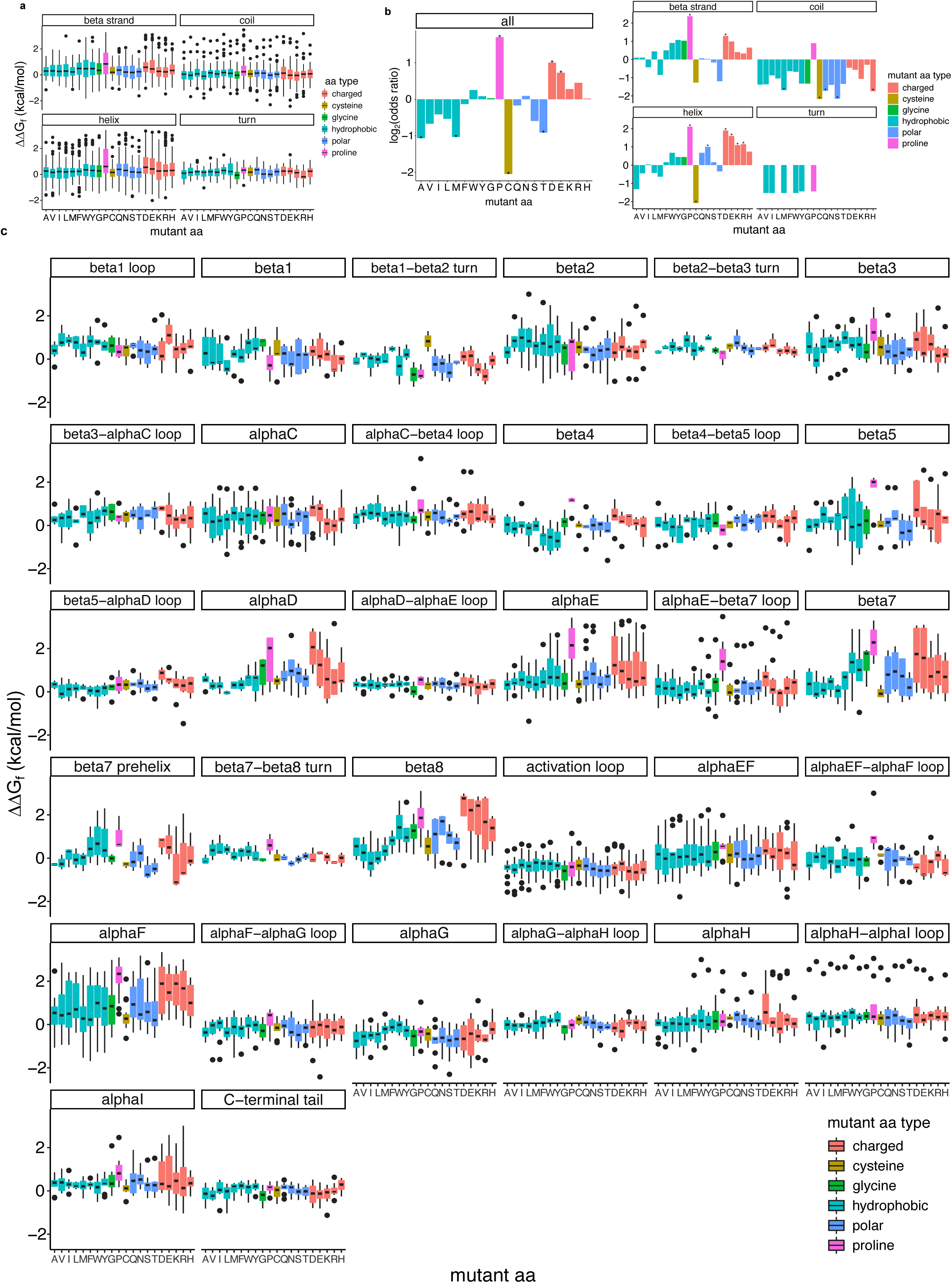
Amino acid preferences for Src kinase domain stability. **a.** Distributions of folding ΔΔG (ΔΔG_f_) in different secondary structure element types according to the identity of the introduced amino acid (‘mutations to’). **b**. Enrichment of mutations according to the introduced amino acid (‘mutations to’) in the destabilizing set across the kinase domain (left panel) and as a function of secondary structure element type (right panel). Significantly enriched or depleted amino acids are labeled with a star (FDR<0.1). **c**. Distributions of ΔΔG_f_ according to the identity of the introduced amino acid (‘mutations to’) in specific secondary structure elements of the Src kinase.

**Supplementary Figure 3:**
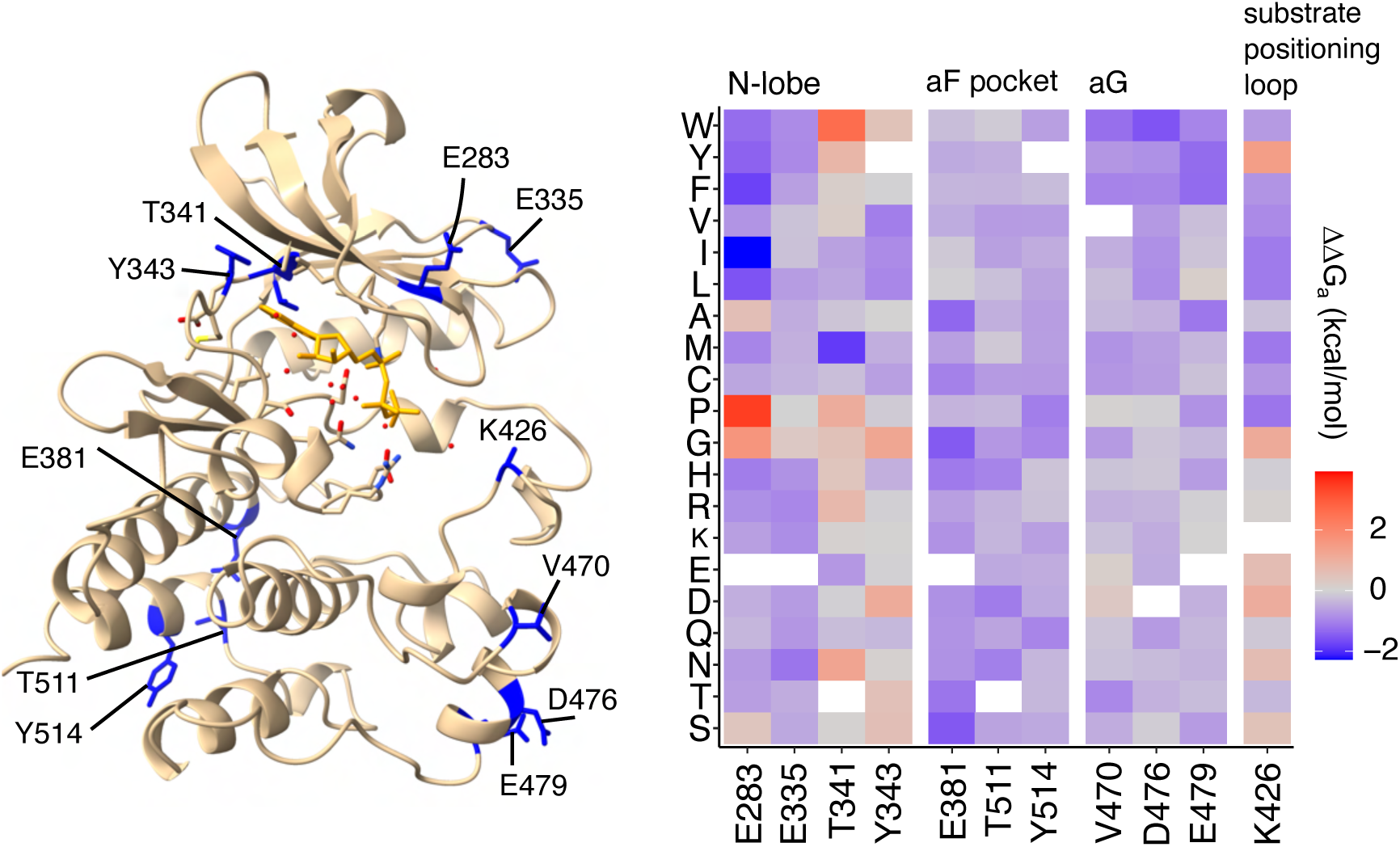
Major activating allosteric sites. Src structure depicting major activatory allosteric sites and heatmap showing ΔΔG_a_ at these sites.

**Supplementary Figure 4:**
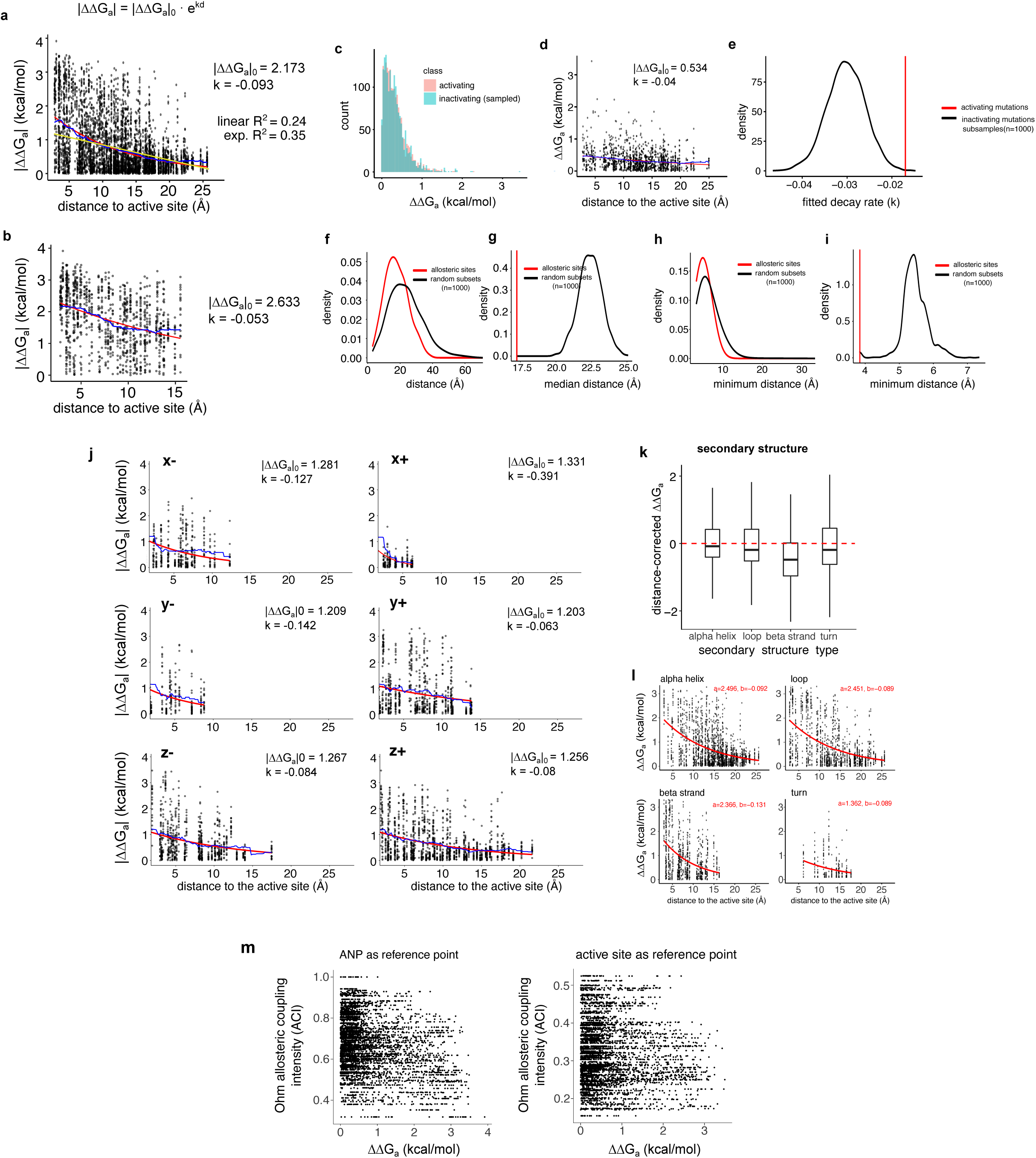
Anisotropy in allosteric transmission across the Src KD. **a.** Exponential decay fits to the relationship between |ΔΔG_a_| to the minimum heavy atom distance to the active site, for all mutations in all Src sites. |ΔΔG_a_|_0_=starting ΔΔG_a_ at distance=0 (active site), k=decay rate, d=distance from the active site. Red line = exponential fit, yellow line = linear model fit, blue = running mean over a 5Å window. **b**. Exponential decay fits to the relationship between |ΔΔG_a_| to the minimum heavy atom distance to the active site, for all mutations in Src allosteric sites and the active site. **c**. Illustrative example of a subsample of inactivating mutations (ΔΔG_a_>0) matching the distribution of effect sizes of activating mutations (ΔΔG_a_<0). **d**. Exponential decay fit to the subsample of inactivating mutations in c. **e**. Distribution of exponential decay rates (k) of the 10,000 subsamples of inactivating mutations (black), compared to the observed decay rate of activating mutations (red line). **f**. Distribution of pairwise distances between Src allosteric sites (red), compared to a null distribution calculated from random subsets of residues (n=1000, black). **g**. Distribution of median pairwise distances calculated from random subsets of residues (n=1000, black), compared to the median pairwise distance between Src allosteric sites (red vertical line). **h**. Distribution of per-site minimum distances to any other allosteric site (red), compared to random subsets of residues (n=1000, black). **i**. Median per-site minimum distance between allosteric sites (red vertical line), compared to the distribution of medians of random subsets (n=1000, black). **j**. Exponential decay fits of |ΔΔG_a_| with distance to the active site in the three orthogonal spatial directions, positive and negative. Decay was calculated in each spatial direction for residues located at a distance less than 10Å from the active site in the two remaining directions. **k**. Distance-corrected ΔΔG_a_ (residuals to loess smoothing curve fit, see Methods) in different secondary structure types. **l**. Exponential decay fits to the relationship between |ΔΔG_a_| to the minimum heavy atom distance to the active site in different secondary structure types.

**Supplementary Figure 5:**
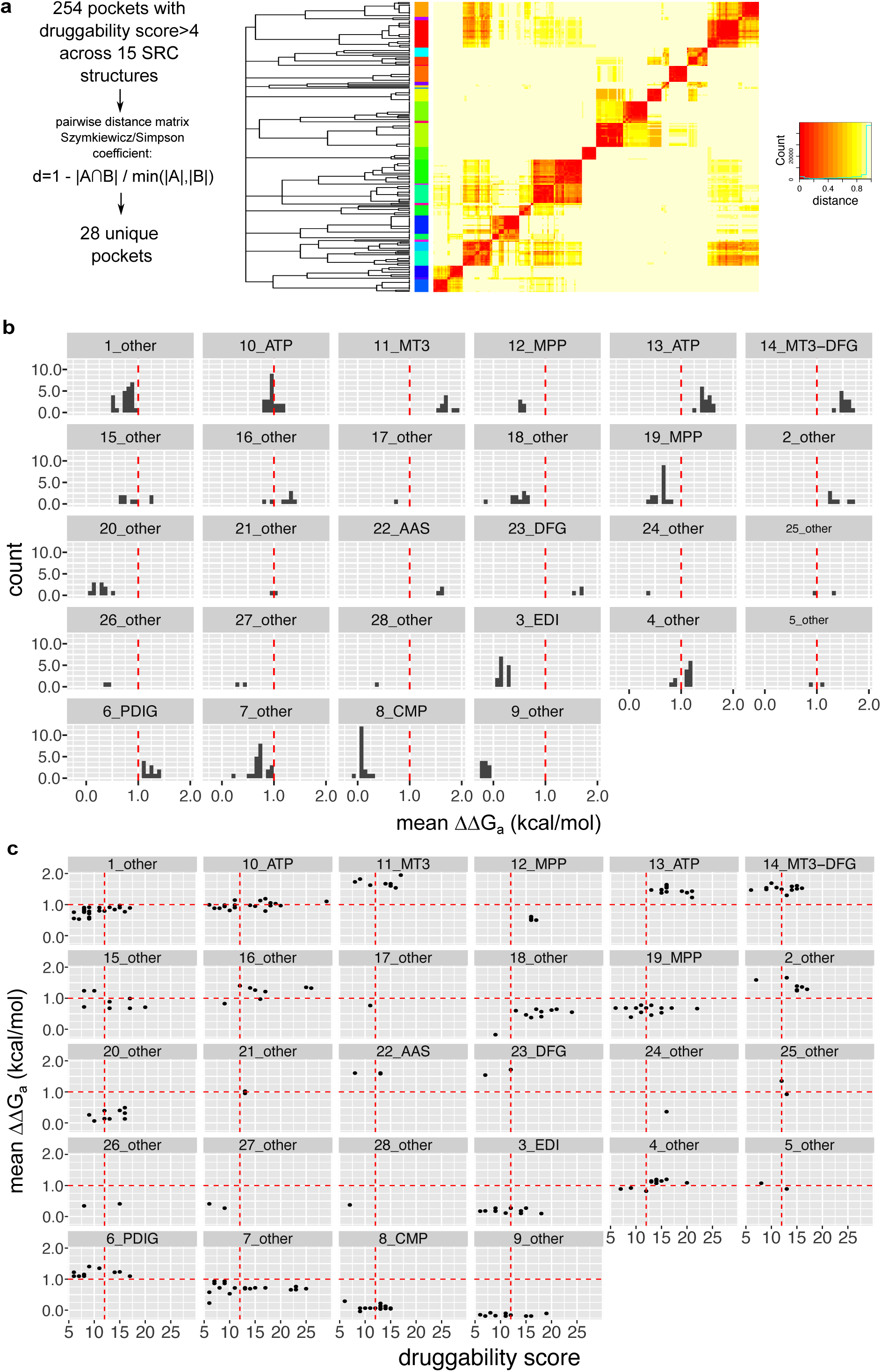
Summary and clustering of Kinase Atlas Src surface pockets. **a**. Clustered matrix of Src surface pockets (defined as residues with minimum heavy atom distance< 5 Å to docked small molecules) across all structures based on the Szymkiewicz-Simpson overlap coefficient. Cluster assignments are shown as a color bar in the left side of the heatmap. **b**. Distribution of mean ΔΔG_a_ of pockets across structures for each of the 28 Src surface pockets. Each data point represents the pocket in a specific Src structure, where present. **c**. Comparison of druggability against mean ΔΔG_a_ across structures for each of the 28 Src surface pockets. Each data point represents the pocket in a specific Src structure, where present.

**Supplementary Figure 6:**
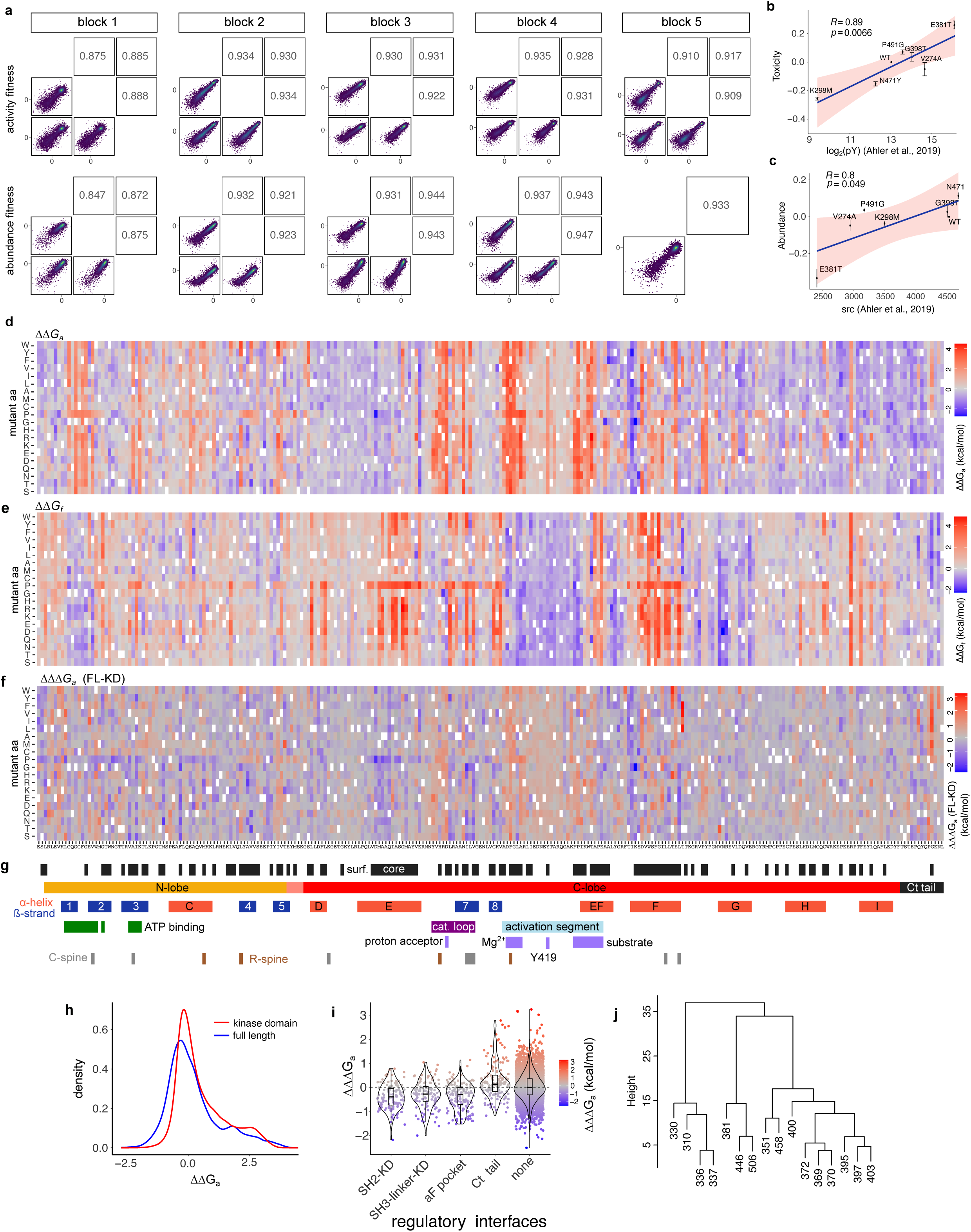
Comparison of full-length and kinase domain alone Src activity energies. **a.** Fitness score replicate correlations for each of the 5 blocks of the Src full-length library, for the activity-dependent toxicity assay (top row) and the sandwich abundancePCA assay (bottom row). **b**. Correlation of activity fitness measurements to in vivo phosphotyrosine levels^43^. **c**. Correlation of sandwich abundancePCA fitness measurements to in vivo Src levels^43^. **d,e**. Heatmaps showing MoCHI inferred changes in activity free energies (d, ΔΔG_a_) and folding free energies (e, ΔΔGf). **f**. Heatmaps showing the changes in ΔΔG_a_ between full-length and kinase domain Src (ΔΔΔG_a_). **g**, Sequence and annotation of Src. Locations of individual secondary structure elements and functional regions were obtained from ^48,71^. **h**. Distributions of ΔΔG_a_ in full-length and kinase domain Src. **i**. Distribution of changes in ΔΔG_a_ between full-length and kinase domain Src (ΔΔΔG_a_) at regulatory domain interfaces, inferred exchanging the full-length and kinase domain alone abundance datasets as the underlying folding data for MoCHI fitting. **j**. Hierarchical clustering of sites with three or more mutations with significantly more activating effects in full-length Src than in the kinase domain alone (residuals to fit>1, FDR<0.1) according to their spatial distances.

**Supplementary Figure 7:**
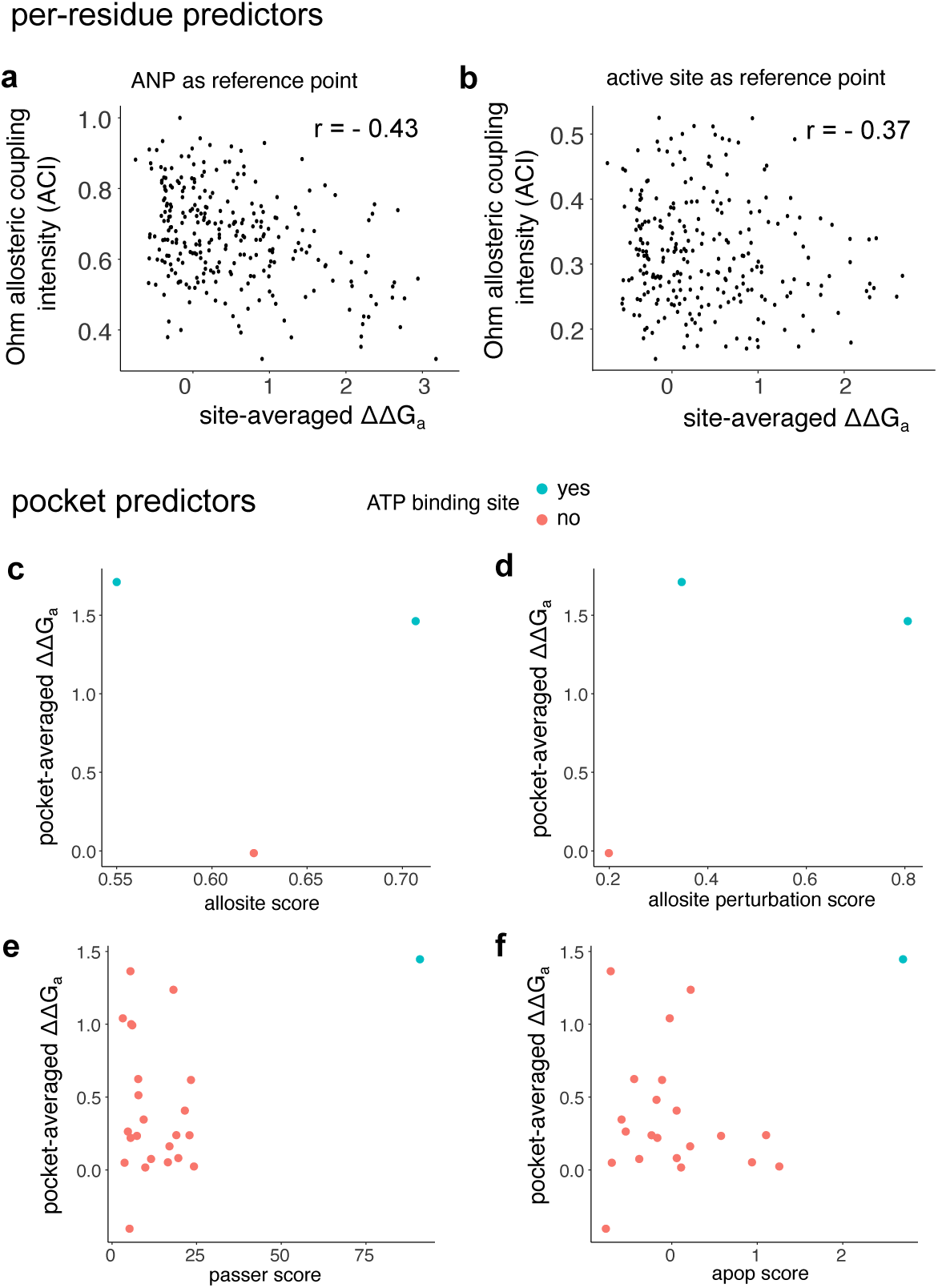
Comparison of allosteric site predictors to Src DMS data. **a-b.** Correlation between site-averaged ΔΔG_a_ and allosteric coupling intensities (ACI) predicted by Ohm^63^ on a residue-level, for full-length Src (a), and for the Src kinase domain alone (b) using 2SRC as reference structure. **c-d**. Comparison of allositePro^63,64^ overall score (c) and perturbation score (d) with pocket-averaged ΔΔG_a_. **e**. Comparison of PASSer^65^ scores with pocket-averaged ΔΔG_a_. **f**. Comparison of apop^66^ scores with pocket-averaged ΔΔG_a_.

## Data availability

All DNA sequencing data have been deposited in the Gene Expression Omnibus under the accession number GSE247740. All scaled fitness measurements and their associated errors for the activity dependent toxicity assay and aPCA are available as Supplementary Table 1. All fitted ΔΔG_f_ and ΔΔG_a_ values and their associated errors are available as Supplementary Table 2. A summary of all Src surface pockets with their druggability scores and ΔΔG_a_ are available as Supplementary Table 3.

## Code availability

Source code used to perform all analyses and to reproduce all figures in this work is available at: https://github.com/lehner-lab/src_allostery. Files required to reproduce the analyses can be downloaded at <u>10.5281/zenodo.10158641</u>.

## Author contributions

A.B. performed all experiments and analyses. A.J.F. helped with thermodynamic model fitting and related analyses. A.B. and B.L. conceived the project, designed analyses, and wrote the manuscript with input from A.J.F.

## Acknowledgements

This work was funded by a European Research Council (ERC) Advanced (883742) grant, the Spanish Ministry of Science and Innovation (LCF/PR/HR21/52410004, EMBL Partnership, Severo Ochoa Centre of Excellence), the Bettencourt Schueller Foundation, the AXA Research Fund, Agència de Gestió d’Ajuts Universitaris i de Recerca (AGAUR, 2017 SGR 1322), the CERCA Program/Generalitat de Catalunya, and Wellcome (Grant reference: 220540/Z/20/A, ‘Wellcome Sanger Institute Quinquennial Review 2021-2026’). A.B was funded by an EMBO (ALTF 183-2020) and Marie Skłodowska-Curie (101030961) fellowship. We thank all members of the Lehner Lab for helpful discussions and suggestions.

## Materials and methods

### Media

- LB: 10 g/L Bacto-tryptone, 5 g/L Yeast extract, 10 g/L NaCl. Autoclaved 20 min at 120°C.
- YPDA: 20 g/L glucose, 20 g/L Peptone, 10 g/L Yeast extract, 40 mg/L adenine sulphate. Autoclaved 20 min at 120°C.
- SORB: 1 M sorbitol, 100 mM LiOAc, 10 mM Tris pH 8.0, 1 mM EDTA.
- Filter sterilized (0.2 mm Nylon membrane, ThermoScientific).
- Plate mixture: 40% PEG3350, 100 mM LiOAc, 10 mM Tris-HCl pH 8.0, 1 mM EDTA pH 8.0. Filter sterilized.
- Recovery medium: YPD (20 g/L glucose, 20 g/L Peptone, 10 g/L Yeast extract) +0.5 M sorbitol. Filter sterilized.
- SC -URA: 6.7 g/L Yeast Nitrogen base without amino acid, 20 g/L glucose, 0.77 g/L complete supplement mixture drop-out without uracil. Filter sterilized.
- SC -URA/ADE: 6.7 g/L Yeast Nitrogen base without amino acid, 20 g/L glucose, 0.76 g/L complete supplement mixture drop-out without uracil, adenine and methionine. Filter sterilized.
- MTX competition medium: SD –URA/ADE + 200 ug/mL methotrexate (BioShop Canada Inc., Canada), 2% DMSO.
- SC -URA 2% Raffinose 0.1% glucose: 6.7 g/L Yeast Nitrogen base without amino acid, 20 g/L raffinose, 1 g/L glucose, 0.77 g/L complete supplement mixture drop-out without uracil. Filter sterilized.
- SC -URA 2% Galactose 0.1% glucose: 6.7 g/L Yeast Nitrogen base without amino acid, 20 g/L galactose, 1 g/L glucose, 0.77 g/L complete supplement mixture drop-out without uracil. Filter sterilized.
- DNA extraction buffer: 2% Triton-X, 1% SDS, 100mM NaCl, 10mM Tris-HCl pH8, 1mM EDTA pH8.

### Plasmid construction

All the Src constructs for expression in yeast and plasmid sequences have been verified by whole plasmid sequencing (Plasmidsaurus). Their sequences and benchling links can be found in Supplementary Table 4. All oligonucleotide sequences used for plasmid construction can be found in Supplementary Table 5. Full length and KD Src sequences used can be found in Supplementary Table 6.

To assay in vivo soluble expression of the Src KD we used pGJJ133, a modified version of the pGJJ045 abundancePCA plasmid where stop codons have been introduced downstream of the NheI-HindIII cloning sites to allow flexible cloning of ORFs that do not contain STOP codons. To assay activity-dependent toxicity of Src, we used pTB022, a plasmid based on the same backbone as the aPCA plasmids but containing a yeast GAL promoter to drive the expression of NheI-HindIII inserts not fused to any DHFR fragment or linker. We ordered a gene block of Src codon-optimized for yeast expression and flanked by NheI-HindIII restriction sites (IDT). We used oligonucleotides oTB063 and oTB064 to amplify the Src KD and introduce an NheI site at the 5′end of the KD sequence, and we cloned the KD fragment on pGJJ133 via restriction digestion and T4 ligation with NheI and HindIII (NEB), resulting in pTB109. The KD fragment was cloned on pTB022 via restriction digestion with NheI and HindIII (NEB) and T4 ligation (NEB), and the resulting plasmid was subjected to a round of site directed mutagenesis (NEB) with oTB215 and oTB216 to introduce a start codon at the beginning of the KD sequence, resulting in pTB112.

To assay in vivo soluble expression of full-length Src, we generated pTB043 via Gibson assembly. pTB043 is based on the same backbone as the aPCA plasmids, and contains a construct where full length Src is fused to the DHFR3 fragment in its N-terminus, and to the DHFR1,2 fragment in its C-terminus. This fusion construct is driven by a cyc promoter and terminated by a cyc terminator, as in the rest of aPCA plasmids. To assay activity-dependent toxicity of full-length Src, we cloned the Src gene block into pTB022 using NheI-HindIII restriction digestion and T4 ligation, resulting in pTB023.

### Variant library design and cloning

To fully cover the Src kinase domain from E268 to L536, the library was divided in 5 overlapping blocks or tiles of ∼60 aa to be cloned and selected separately. Within each tile, we chose 10 genetic backgrounds with a wide range of effects on Src kinase activity based on a previous deep mutagenesis dataset^43^. The sequences of all tiles, constant regions, and genetic backgrounds are in Supplementary Table 4.

The library was ordered as two IDT oPools (Pool 1 with block1 and Pool 2 with blocks 2-5), containing all NNK single mutants in each of the 10 backgrounds of each block. 2.5 ul 0.1 uM oPool material was used as a template in a 100 ul Q5 high-fidelity 10 cycle PCR reaction. Primers specific to the constant regions of each block were used (see oligonucleotides). The amplified products were verified on an agarose gel, and column-purified (QIAquick PCR purification kit, QIAGEN) for Gibson assembly. The library was assembled on pTB112 (KD) and on pTB023 (full-length). To do so, the plasmids were linearized with primers pointing outwards from the constant regions of each block (bb primers, see oligonucleotides) so that each linearized vector had at least 20 nt of homology to the amplified oligo pool containing the variants. 120 ng of linearized plasmid were assembled with the amplified oligo pools in a 20 ul Gibson reaction using in-house prepared Gibson assembly mix, and incubating for 3h at 50C. The reaction products were dialyzed using 0.025 uM MCE membranes, concentrated to 5 ul using a SpeedVac machine, and transformed into NEB 10β High-efficiency Electrocompetent *E. coli* cells according to the manufacturer’s protocol. Cells were left to recover in SOC medium (NEB 10β Stable Outgrowth Medium) for 30 minutes, a 2 ul aliquot was plated to quantify the total number of clones, and the rest of the cell volume was transferred to 100 mL of LB medium with ampicillin overnight. 100 mL of each saturated *E. coli* culture were harvested next morning to extract the plasmid library using the Plasmid Plus Midi Kit (QIAGEN). The assemblies were verified using Sanger sequencing (Eurofins). The assembled mutated Src library constructs were transferred into the pGJJ133 and pTB198 plasmids using NheI-HindIII digestion (NEB) and overnight temperature cycling T4 ligation (NEB), followed by dialysis, electroporation, overnight LB-ampicillin selection, and plasmid isolation as described above.

### Large-scale transformation and competition assays

Each library corresponding to each of the 5 blocks was transformed in triplicate, and with a coverage of ∼100x or greater. Three 500 mL YPDA cultures of late log phase *S. cerevisiae* BY4741 cells (OD∼0.8-1) were harvested in 50 mL Falcon tubes, each resuspended in 22 mL SORB medium and incubated for 30 min on a shaker at room temperature. 437.5 ul 10 mg/mL previously boiled (5 min, 100C) ssDNA was added to the cells, and the mix was separated in 5 aliquots of 4.3 mL in 50 mL Falcon tubes, one for each library block. 3 ug library plasmid DNA was added to each aliquot, followed by 17.5 mL Plate Mixture, and the mix was incubated for 30 min on a shaker at room temperature. 1.75 mL DMSO were then added, and the cells were incubated at 42C for 20 min. Following incubation, cells were centrifuged for 3 min at 3,000g, the supernatant was discarded with a pump, and cells were resuspended in Recovery Media and incubated at 30C for 1h. Cells were then centrifuged for 3 min at 3,000g and transferred into 100 mL SC –URA. 10 ul of this culture were immediately plated onto SC -URA selective plates to monitor transformation efficiency. The rest of the culture was incubated overnight at 30C.

For the activity-dependent toxicity assays, the overnight SC -URA cultures were used to inoculate the next day a 100 mL culture of SC -URA with 2% raffinose and 0.1% glucose at an OD=0.2-0.4, which was grown overnight. Cells from this culture were inoculated the next day in 100 mL SC -URA with 2% galactose and 0.1% glucose at an OD=0.05, to induce overexpression of the Src variant library. The remaining input cells grown in 2% raffinose 0.1% glucose were harvested and frozen for DNA extraction (inputs). The galactose cultures were left to grow overnight to an OD=1.6-2.5, corresponding approximately to 5 generations, harvested, and frozen for DNA extraction (outputs).

For the aPCA and sandwichPCA assays, the overnight SC -URA cultures were used to inoculate the next day a 100 mL culture of SC -URA-ADE at an OD=0.2-0.4, which was grown overnight (input culture). Cells from this culture were inoculated in 100 mL SC-URA/ADE +200 ug/ml MTX to select stably expressed Src variants. The remaining input cells grown SC -URA/ADE were harvested and frozen for DNA extraction. The MTX cultures were left to grow overnight to an OD=1.6-2.5, corresponding approximately to 5 generations, harvested, and frozen for DNA extraction (outputs).

### DNA extraction, plasmid quantification and sequencing library preparation

Total DNA was extracted from yeast pellets equivalent to 50 mL of cells at OD=1.6 as described in our previous work^12,30^. Plasmid concentrations in the resulting samples were quantified by against a standard curve of known concentrations by qPCR, using oGJJ152 and oGJJ153 as qPCR primers that amplify in the origin of replication of both toxicity and aPCAa assay plasmids.

To generate the sequencing libraries, we performed two rounds of PCR amplification. In the first round, we used primers flanking the mutated regions in each sample (5 pairs of PCR1 primers, one for each of the Src blocks). This PCR1 reaction allows increasing the nucleotide complexity of the first sequenced bases by introducing frame-shift bases between the Illumina adapters and the sequencing region of interest. For block 5, a different reverse frameshifting pool was used for the sandwichPCA libraries as they differ in the region downstream of the STOP codon of Src (oTB470+ was used instead of oGJJ589+). All frameshifting PCR1 primers are listed in the Oligonucleotides table. 125 million plasmid molecules were used as templates and were amplified for 8 cycles. The reactions were column-purified (QIAquick PCR purification kit, QIAGEN), and 40 ng DNA were used as template for a PCR2 reaction with the standard i5 and i7 primers to add the remainder of the Illumina adapter sequences and the demultiplexing indices (dual indexing) unique to each sample. This PCR2 was run for 8 cycles, and the resulting amplicons were run on a 2% agarose gel to quantify and pool the samples for joint purification, and to ensure the specificity of the amplification and check for any potential excess amplification problems.

The final libraries were size selected by electrophoresis on a 2% agarose gel, and gel-purified (QIAEX II Gel Extraction Kit ,QIAGEN). The amplicons were subjected to Illumina paired end 2×150 sequencing on a NextSeq2000 instrument at the CRG Genomics facility.

### Sequencing data processing and thermodynamic modeling

FastQ files from paired-end sequencing of all aPCA and toxicity experiments were processed with DiMSum^72^ v1.2.11 (https://github.com/lehner-lab/DiMSum) using default settings with minor adjustments. A minimum input read count threshold was set for 1-nucleotide substitutions using the “fitnessMinInputCountAny” option, in order to minimize the fraction of reads per variant related to sequencing error-induced “variant flow” from the WT. The option “barcodeIdentityPath” was used to specify a variants file in order to recover only the variants corresponding to the designed library (NNK mutations in one of the predefined genetic backgrounds).

We used MoCHI^31^ (https://github.com/lehner-lab/MoCHI) to fit two global mechanistic models, one for the Src KD and one for full-length Src, using the corresponding 10 aPCA and toxicity assay datasets (2 molecular phenotypes x 5 blocks) simultaneously. The software is based on our previously described genotype-phenotype modeling approach^12^ with additional functionality and improvements for ease-of-use and flexibility^30,31^.

We fit a phenomenological ‘enzyme folding and activation’ model to the Src kinase data. To do so, we use an explicit three-state thermodynamic model in which the protein can exist in three states: unfolded and inactive (ui), folded and inactive (fi), and folded and active (fa). The folding energy, ΔGf, is defined as the energy difference between the unfolded (ui) and folded (fi) inactive states, and the ‘activity’ energy, ΔG_a_, is defined as the energy difference between the inactive (fi) and active (fa) folded states. We note that this ΔG_a_ parameter quantifies all changes in the activity of Src not explained by changes in abundance, including changes in the equilibrium between active and inactive conformations of the enzyme, but also any other effects on catalysis (kcat) or substrate specificity (km) not related to conformational changes. We assume that the probability of the unfolded and active state (ua) is negligible, and that changes in folding and active state energies are additive i.e. the total free energy change of an arbitrary variant relative to the wild-type sequence is simply the sum over residue-specific energy changes corresponding to all constituent single amino acid substitutions.

We set MoCHI parameters to specify a neural network architecture consisting of two additive trait layers, one for folding and one for active state energies, as well as one linear transformation layer per experiment (5x for toxicity and 5x for aPCA blocks). The specified non-linear transformations “TwoStateFractionFolded” and “ThreeStateFractionBound” derived from the Boltzmann distribution function relate energies to proportions of folded molecules, and molecules in the active vs inactive states of Src respectively. We used as input data the WT, single, and double aa variants of Src (--order_subset 0,1,2) and modeled the data in the absence of interactions between single substitutions (--max_interaction_order 1). DiMSum output fitness tables were formatted to include the full 5-block sequence, and the sign of the toxicity assay fitness was reversed as Src activity and cellular fitness are anti-correlated. We additionally removed variants corresponding to the activity dead and unstable L494P genetic background as the fitness range of this background is extremely narrow and we find it greatly biased block 5 fitted energies towards negative values. The training and validation workflow, as well as the confidence estimation used was as described in our previous work^12,30^ but without L2 regularization as we did not observe artifactually large values of fitted free energy changes. For model performance estimates, the maximum explainable variance (of genetic origin) was calculated by subtracting the total estimated technical variance (as reported by DiMSum fitness errors) from the total fitness variance i.e. 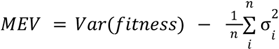 where σ*_i_* is is the fitness error associated with variant *i* and *n* is the total number of variants. The fraction of explainable variance is then given by *FEV* = *MEV*/*Var*(*fitness*). In order to minimize the effect of outlier fitness errors on FEV estimates, we discarded variants whose errors were in the top 15th percentile. We evaluated performance on the retained variants and computed the explainable variance captured by MoCHI models as the predicted versus observed R^2^ on held out double mutants divided by the FEV.

We additionally fit a 2-state model following the same procedure as above but where mutant effects on both activity and abundance fitness are assumed to be driven by a single underlying biophysical trait (“TwoStateFractionFolded”), and a 4-state model in which mutations can have independent effects on two distinct active state conformations (“FourStateFractionBound”).

### Identification of activating, inactivating, stabilizing and destabilizing mutations, and enrichments in secondary structure elements or functional regions

The mean weights (‘mean kcal/mol’) and standard deviations (‘sd kcal/mol’) from MoCHI fits were used for statistical testing to identify mutations with changes in stability or activity, using z-tests, where *Z* = (*ref*. *value* − *mean*)/*sd*. P-values were calculated on the basis of a normal distribution. Enrichments of particular mutation classes in individual sites, or in subgroups of residues based on structural or functional annotations were tested using Fisher’s exact test and comparing to the rest of the KD as background unless specified otherwise. In all box plots shown throughout the manuscript, the central line represents the median, the lower and upper hinges correspond to the first and third quartiles, the upper whisker extends from the hinge to the largest value no further than 1.5 * IQR (inter-quartile range) from the hinge, and the lower whisker extends from the hinge to the smallest value at most 1.5 * IQR of the hinge.

### Quantification of the distance dependence of mutation effects

To quantify the dependence of mutation effects on the distance from the active site, we computed the minimum distance between all atoms of each residue of Src and the active site using the 2SRC^67^ structure. We computed distances to (1) the nucleotide (the non-hydrolyzable ATP analog AMP-PNP in 2SRC), and (2) the catalytic D388, and (3) the proposed phosphosite substrate positioning residue P428^48^, and took the minimum distance for each residue to these 3 reference points. To fit an exponential curve to the data, we used the R stats package. We first used the optim() function to select reasonable starting values, to then fit the *y* = *a* · *e*^*bx*^ curve using the nls() function, where a is the estimate of |ΔΔG | at the active site (|ΔΔG_a_|_0_), b is the decay rate (k), and x is the minimum heavy atom residue distance to the active site. To estimate distance-corrected mutation effects, we used the msir package^73^ to fit a loess smoothing curve and quantified the residuals to the fit across different secondary structure element types. To quantify allosteric decay rate variation across different spatial directions, we used the x,y,z directions as defined in the 2SRC pdb entry. To quantify decay in each direction, we considered residues at a distance of 10Å or less from the active site in the two remaining orthogonal directions.

To compare the distance dependence of activating and inactivating mutations independently of their effect sizes, we subsampled the set of mutations with positive ΔΔG_a_ to match the number and distribution of effects of those with negative ΔΔG_a_ (n=10,000 subsamples, with replacement), and fit exponential decays to these. A p-value was calculated as the fraction of subsamples in which the decay is lower or equal than that of activating mutations.

To test for clustering of allosteric sites, we computed the distribution of C_α_-C_α_ pairwise distances between all allosteric sites in the Src KD, and compared it to a null distribution obtained from 1000 subsets of randomly sampled residues of equal number. A p-value was calculated as the fraction of subsets in which the median distance is lower or equal to the observed median distance. To test for connectivity, we took a similar approach but computing the distribution of minimum distances from each allosteric site to any other allosteric site.

### Quantification of structural features and contacts

The locations of individual secondary structure elements and functional annotations were obtained from ^48,71^. Solvent accessible surface area were calculated using freeSASA v2.0.3^74^ with parameters -n 20 --format=rsa --radii=naccess, and with 2SRC as a reference structure, both using the full length Src structure, and using the KD residues alone. Secondary structure type annotations (helix, beta sheet, turn, loop) from uniprot were used.

To define active and inactive state contacts, we used getcontacts (https://getcontacts.github.io/) on representative structures of the inactive (2SRC) and active (1Y57^57^) states. Prior to defining contacts we added hydrogen atoms to the structures using the pymol h_add method. We then used get_static_contacts.py with parameters --itypes all.

For the following analyses we considered only salt bridges, pi-cation interactions, side chain-side chain hydrogen bonds, and side chain-backbone hydrogen bonds as the rest of contact types did not display conformational state specificity. Contacts of the same type and between the same residues were collapsed into a single contact, and duplicated contacts annotated both as salt bridge and side chain-side chain hydrogen bond were collapsed as salt bridge. We defined four types of residues based on their contact patterns: active residues engaged in contacts exclusively in the active state, inactive residues engaged in contacts only in the inactive state, non-specific residues engaged in the same contact in both structures, and swapping residues that engage in different contacts in the active and inactive conformations. To display the contacts in Src structures, we used a pseudobond representation in ChimeraX^75^, with the thickness of the contact being proportional to the averaged mutation effect of the two contacting residues. Contacts between swapping residues were represented as dashed pseudobonds.

### Allostery prediction

We fit linear models to predict ΔΔG_a_ using the base R lm() function, using as predictors the wt and mutant aa, secondary structure type in which the mutation is located, the specific secondary structure element in which the mutation is located, rSASA, contact type, and the residue type classification (active, inactive, swapping, both, or none) according to their contact patterns as described above. We also used as predictors the distance to D389 (catalytic site), and to the nucleotide (AMP-PNP in 2SRC), transformed according to the fitted exponential decay: *d_tr = e_*^*k*\**d*^ where d_tr_ is the transformed distance, k = - 0.093 Å^-1^, and d is the distance.

Models were evaluated on held out data using a 10-fold cross validation strategy. To evaluate the performance of allosteric site prediction models on predicting the Src dataset, we generated allosteric pocket predictions with allositePro^64^, PASSer^65^, APOP^66^, and compared the pocket scores of these predictors to the mean averaged ΔΔG_a_ of all constituent residues. In addition, we generated residue-level allosteric coupling intensity predictions (ACI) with Ohm^63^ and compared them directly to ΔΔG_a_.

### Analysis of Src surface pockets

We used the Kinase Atlas^25^ to retrieve all possible Src potentially druggable surface pockets. We used the docking analyses of all 15 available Src structures, and we defined each potential Src surface pocket as the set of residues located at a minimum distance of 5 Å from a cluster of docked molecules, resulting in a total of 384 pockets distributed across the 15 structures. After filtering out pockets with a druggability score <5, we collapsed the remaining 254 pockets into a final set of unique pockets as many are present in multiple structures. To do so, we calculated a pairwise distance matrix between all pockets, using as a distance metric 1 minus the Szymkiewicz-Simpson overlap coefficient. We applied hierarchical clustering to the distance matrix, resulting in 28 unique pockets.

As each of these 28 unique pockets is a set of pockets present across several structures, we summarized the total number of structures in which each is found, and calculated the average and maximum druggability across all structures. For each pocket in each structure, we averaged the ΔΔG_a_ per residue, and calculated the mean residue-averaged ΔΔG_a_ (mean ΔΔG_a_), maximum residue-averaged ΔΔG_a_ (max ΔΔG_a_), and minimum ΔΔG_a_ (min ΔΔG_a_). We also calculated the odds ratio of enrichment of each pocket in activating and inactivating mutations relative to the rest of the KD, and tested its statistical significance using Fisher’s exact test. As these metrics were very consistent across structures, we used the median across structures as the final unique pocket summary shown in Figure 5b.

### Quantification of changes in mutational effects by regulatory domains

To quantify the changes in ΔΔG_a_ that result from the presence of Src regulatory domains, we fit a linear model to the ΔΔG_a_ dataset in full length Src against ΔΔG_a_ in the kinase domain alone, and used the residuals to the fit as an estimator of ΔΔΔG_a_ (full length minus kinase domain). To identify mutations with significant ΔΔΔG_a_, we calculated z-scores incorporating the errors of both full-length (sdFL) and kinase domain alone (sdKD) ΔΔG_a_, as 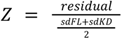. We defined interdomain interfaces as residues involved in direct contacts between the kinase domain and the regulatory domains (SH3, SH2, linker) using getcontacts, and excluding water bridges. To identify spatial clusters, we selected sites with at least 2 mutations with ΔΔΔG_a_< −1 and FDR<0.1 and applied hierarchical clustering to a matrix with their pairwise C_α_-C_α_ distances.

