## Supplementary material for "The allosteric landscape of the Src kinase"

### Supplementary tables

**Supplementary table 1:** Fitness scores and errors
**Supplementary table 2:** Fitted values of ΔΔGf and ΔΔGa
**Supplementary table 3:** Src surface pocket summary

**Supplementary table 4:** Plasmids

| | | |
|---|---|---|
| pGJJ133 | aPCA empty | Available upon request (Material Transfer Agreement required) |
| pTB022 | toxicity empty | https://benchling.com/s/seq-rGJbDMKRao9IdlAqhMoJ?m=slm-cUTBKgetTLCpJ3TveV5w |
| pTB198 | sandwichPCA empty | https://benchling.com/s/seq-mve4HVJwo23HnO6BPNym?m=slm-lNfOeXfnOvLHoarUjhPK |
| pTB109 | aPCA Src KD | https://benchling.com/s/seq-obS1wabcFstnPlwNCdPI?m=slm-an1b6yZtzRU1xIytarI5 |
| pTB112 | toxicity Src KD | https://benchling.com/s/seq-PUIubmPUQrNsuv9YkjJM?m=slm-KBT9Fq1il6zXODI1mvnj |
| pTB043 | sandwichPCA full length | https://benchling.com/s/seq-0sA5ySViNYIlnltmMAFI?m=slm-27zYqdcFKogMjbIGqR1y |
| pTB023 | toxicity full length | https://benchling.com/s/seq-PdGjC1KRegS3IUGnMNPh?m=slm-CCYco9fv12T7nVaTxXhf |

**Supplementary table 5:** Oligonucleotides

| | |
|---|---|
| Amplification of SRC KD from full length gBlock | |
| oTB063 | CAATATGCTAGCGATGCTTGGGAGATCCCTC |
| oTB064 | TAATTTAAAGCTTCAAGTTCTCTC |
| Introduction of start codon in pTB112 | |
| oTB214 | atgGATGCTTGGGAGATCCCTC |
| oTB215 | GCTAGCCTCCTTGACGTT |
| oPool and backbone amplification for Gibson assembly | |
| oTB447_b1_ins_F | GATGCTTGGGAGATCCCTC |
| oTB448_b1_ins_R | CACTCACAACTGCATACAATTG |
| oTB449_b2_ins_F | AGCACAAGTCATGAAGAAGC |
| oTB450_b2_ins_R | CACGGTGTACGTAATTCATTC |
| oTB451_b3_ins_F | CATGGCCGCCCAGATTG |
| oTB452_b3_ins_R | CGTGAACCTTCCATATAAGGC |
| oTB453_b4_ins_F | GTGCAAAGTTCCCCATCAAG |
| oTB454_b4_ins_R | CATAAGGTCGTGCAAGCTC |
| oTB455_b5_ins_F | GAACGTGGTTATAGAATGCC |

| | |
|---|---|
| oTB220_b5_ins_R | GCGTGACATAACTAATTTAAAGC |
| oTB457_b1_bb_F | CAATTGTATGCAGTTGTGAGTG |
| oTB458_b1_bb_R | GAGGGATCTCCCAAGCATC |
| oTB459_b2_bb_F | GAATGAATTACGTACACCGTG |
| oTB460_b2_bb_R | AGCTTCTTCATGACTTGTGC |
| oTB461_b3_bb_F | GCCTTATATGGAAGGTTCACG |
| oTB462_b3_bb_R | CAATCTGGGCGGCCATG |
| oTB463_b4_bb_F | GAGCTTGCACGACCTTATG |
| oTB464_b4_bb_R | CTTGATGGGGAACTTTGCAC |
| oTB241_b5_bb_F | AAGCTTTAAATTAGTTATGTCACG |
| oTB466_b5_bb_R | GGCATTCTATAACCACGTTC |
| qPCR quantification oligos | |
| oGJJ152 | GCCTACATACCTCGCTCTGC |
| oGJJ153 | CAACCCGGTAAGACACGACT |
| Frameshifting PCR1 oligos | |
| oTB302_b1_fs_F | ACACTCTTTCCCTACACGACGCTCTTCCGATCTATGCTTGGGAGATCCCTC |
| oTB303_302_+1 | ACACTCTTTCCCTACACGACGCTCTTCCGATCTNATGCTTGGGAGATCCCTC |
| oTB304_302_+2 | ACACTCTTTCCCTACACGACGCTCTTCCGATCTNCATGCTTGGGAGATCCCTC |
| oTB305_302_+3 | ACACTCTTTCCCTACACGACGCTCTTCCGATCTGGCATGCTTGGGAGATCCCTC |
| oTB306_302_+4 | ACACTCTTTCCCTACACGACGCTCTTCCGATCTCTGNATGCTTGGGAGATCCCTC |
| oTB307_302_+5 | ACACTCTTTCCCTACACGACGCTCTTCCGATCTNWWANATGCTTGGGAGATCCCTC |
| oTB308_b1_fs_R | GTGACTGGAGTTCAGACGTGTGCTCTTCCGATCTCACTCACAACTGCATACAATTG |
| oTB309_308_+1 | GTGACTGGAGTTCAGACGTGTGCTCTTCCGATCTGCACTCACAACTGCATACAATTG |
| oTB310_308_+2 | GTGACTGGAGTTCAGACGTGTGCTCTTCCGATCTAGCACTCACAACTGCATACAATTG |
| oTB311_308_+3 | GTGACTGGAGTTCAGACGTGTGCTCTTCCGATCTTAGCACTCACAACTGCATACAATTG |
| oTB312_308_+4 | GTGACTGGAGTTCAGACGTGTGCTCTTCCGATCTWTTGCACTCACAACTGCATACAATTG |
| oTB313_308_+5 | GTGACTGGAGTTCAGACGTGTGCTCTTCCGATCTSCATGCACTCACAACTGCATACAATTG |
| oTB471_449_b2_FS_F | ACACTCTTTCCCTACACGACGCTCTTCCGATCTAGCACAAGTCATGAAGAAGC |

| | |
|---|---|
| oTB472_449_+1 | ACACTCTTTCCCTACACGACGCTCTTCCGATCTTAGCACAAGTCATGAAGAAGC |
| oTB473_449_+2 | ACACTCTTTCCCTACACGACGCTCTTCCGATCTCTAGCACAAGTCATGAAGAAGC |
| oTB474_449_+3 | ACACTCTTTCCCTACACGACGCTCTTCCGATCTGCTAGCACAAGTCATGAAGAAGC |
| oTB475_449_+4 | ACACTCTTTCCCTACACGACGCTCTTCCGATCTNNNTAGCACAAGTCATGAAGAAGC |
| oTB476_449+5 | ACACTCTTTCCCTACACGACGCTCTTCCGATCTNNNTTAGCACAAGTCATGAAGAAGC |
| oTB477_450_b2_FS_R | GTGACTGGAGTTCAGACGTGTGCTCTTCCGATCTCACGGTGTACGTAATTCATTC |
| oTB478_450_+1 | GTGACTGGAGTTCAGACGTGTGCTCTTCCGATCTGCACGGTGTACGTAATTCATTC |
| oTB479_450_+2 | GTGACTGGAGTTCAGACGTGTGCTCTTCCGATCTTGCACGGTGTACGTAATTCATTC |
| oTB480_450_+3 | GTGACTGGAGTTCAGACGTGTGCTCTTCCGATCTATGCACGGTGTACGTAATTCATTC |
| oTB481_450_+4 | GTGACTGGAGTTCAGACGTGTGCTCTTCCGATCTWSTTCACGGTGTACGTAATTCATTC |
| oTB482_450_+5 | GTGACTGGAGTTCAGACGTGTGCTCTTCCGATCTSWNWWCACGGTGTACGTAATTCATTC |
| oTB483_451_b3_FS_F | ACACTCTTTCCCTACACGACGCTCTTCCGATCTCATGGCCGCCCAGATTG |
| oTB484_451_+1 | ACACTCTTTCCCTACACGACGCTCTTCCGATCTGCATGGCCGCCCAGATTG |
| oTB485_451_+2 | ACACTCTTTCCCTACACGACGCTCTTCCGATCTAGCATGGCCGCCCAGATTG |
| oTB486_451_+3 | ACACTCTTTCCCTACACGACGCTCTTCCGATCTTTGCATGGCCGCCCAGATTG |
| oTB487_451_+4 | ACACTCTTTCCCTACACGACGCTCTTCCGATCTNNNNCATGGCCGCCCAGATTG |
| oTB488_451_+5 | ACACTCTTTCCCTACACGACGCTCTTCCGATCTNNNNWCATGGCCGCCCAGATTG |
| oTB489_452_b3_FS_R | GTGACTGGAGTTCAGACGTGTGCTCTTCCGATCTCGTGAACCTTCCATATAAGGC |
| oTB490_452_+1 | GTGACTGGAGTTCAGACGTGTGCTCTTCCGATCTACGTGAACCTTCCATATAAGGC |
| oTB491_452_+2 | GTGACTGGAGTTCAGACGTGTGCTCTTCCGATCTGACGTGAACCTTCCATATAAGGC |
| oTB492_452_+3 | GTGACTGGAGTTCAGACGTGTGCTCTTCCGATCTTTACGTGAACCTTCCATATAAGGC |
| oTB493_452_+4 | GTGACTGGAGTTCAGACGTGTGCTCTTCCGATCTWSSACG |

| | |
|---|---|
| | TGAACCTTCCATATAAGGC |
| oTB494_452_+5 | GTGACTGGAGTTCAGACGTGTGCTCTTCCGATCTSWWWNCGTGAACCTTCCATATAAGGC |
| oTB495_453_b4_FS_F | ACACTCTTTCCCTACACGACGCTCTTCCGATCTGTGCAAAGTTCCCCATCAAG |
| oTB496_453_+1 | ACACTCTTTCCCTACACGACGCTCTTCCGATCTAGTGCAAAGTTCCCCATCAAG |
| oTB497_453_+2 | ACACTCTTTCCCTACACGACGCTCTTCCGATCTTAGTGCAAAGTTCCCCATCAAG |
| oTB498_453_+3 | ACACTCTTTCCCTACACGACGCTCTTCCGATCTCCCGTGCAAAGTTCCCCATCAAG |
| oTB499_453_+4 | ACACTCTTTCCCTACACGACGCTCTTCCGATCTNNAAGTGCAAAGTTCCCCATCAAG |
| oTB500_453_+5 | ACACTCTTTCCCTACACGACGCTCTTCCGATCTNNNNNGTGCAAAGTTCCCCATCAAG |
| oTB501_454_b4_FS_R | GTGACTGGAGTTCAGACGTGTGCTCTTCCGATCTCATAAGGTCGTGCAAGCTC |
| oTB502_454_+1 | GTGACTGGAGTTCAGACGTGTGCTCTTCCGATCTGCATAAGGTCGTGCAAGCTC |
| oTB503_454_+2 | GTGACTGGAGTTCAGACGTGTGCTCTTCCGATCTTGCATAAGGTCGTGCAAGCTC |
| oTB504_454_+3 | GTGACTGGAGTTCAGACGTGTGCTCTTCCGATCTATGCATAAGGTCGTGCAAGCTC |
| oTB505_454_+4 | GTGACTGGAGTTCAGACGTGTGCTCTTCCGATCTNNNGCATAAGGTCGTGCAAGCTC |
| oTB506_454_+5 | GTGACTGGAGTTCAGACGTGTGCTCTTCCGATCTNNNSGCATAAGGTCGTGCAAGCTC |
| oTB507_455_b5_FS_F | ACACTCTTTCCCTACACGACGCTCTTCCGATCTGAACGTGGTTATAGAATGCC |
| oTB508_455_+1 | ACACTCTTTCCCTACACGACGCTCTTCCGATCTTGAACGTGGTTATAGAATGCC |
| oTB509_455_+2 | ACACTCTTTCCCTACACGACGCTCTTCCGATCTCTGAACGTGGTTATAGAATGCC |
| oTB510_455_+3 | ACACTCTTTCCCTACACGACGCTCTTCCGATCTACTGAACGTGGTTATAGAATGCC |
| oTB511_455_+4 | ACACTCTTTCCCTACACGACGCTCTTCCGATCTNNCTGAACGTGGTTATAGAATGCC |
| oTB512_455_+5 | ACACTCTTTCCCTACACGACGCTCTTCCGATCTNNNNTGAACGTGGTTATAGAATGCC |
| oGJJ589_b5_R | GTGACTGGAGTTCAGACGTGTGCTCTTCCGATCTGCGTGACATAACTAATTTAAAGC |
| oGJJ590_589_+1 | GTGACTGGAGTTCAGACGTGTGCTCTTCCGATCTNGCGTGACATAACTAATTTAAAGC |

| | |
|---|---|
| oGJJ591_589_+2 | GTGACTGGAGTTCAGACGTGTGCTCTTCCGATCTNNGCGTGACATAACTAATTTAAAGC |
| oGJJ592_589_+3 | GTGACTGGAGTTCAGACGTGTGCTCTTCCGATCTHNHGCGTGACATAACTAATTTAAAGC |
| oGJJ593_589_+4 | GTGACTGGAGTTCAGACGTGTGCTCTTCCGATCTHWWHGCGTGACATAACTAATTTAAAGC |
| oGJJ594_589_+5 | GTGACTGGAGTTCAGACGTGTGCTCTTCCGATCTHWWAAGCGTGACATAACTAATTTAAAGC |
| oTB513_470+ | GTGACTGGAGTTCAGACGTGTGCTCTTCCGATCTCCCACCACCTCCactAAG |
| oTB514_470_+1 | GTGACTGGAGTTCAGACGTGTGCTCTTCCGATCTATCCCACCACCTCCactAAG |
| oTB515_470_+2 | GTGACTGGAGTTCAGACGTGTGCTCTTCCGATCTTGACCCACCACCTCCactAAG |
| oTB516_470_+3 | GTGACTGGAGTTCAGACGTGTGCTCTTCCGATCTGAGTCCCACCACCTCCactAAG |
| oTB517_470_+4 | GTGACTGGAGTTCAGACGTGTGCTCTTCCGATCTSWTGDCCCACCACCTCCactAAG |
| oTB518_470_+5 | GTGACTGGAGTTCAGACGTGTGCTCTTCCGATCTWSNNDCCCACCACCTCCactAAG |

**Supplementary Table 6: Src sequences**

<u>Src kinase domain:</u>
CTAGCGATGCTTGGGAGATCCCTCGTGAATCACTGCGTCTTGAGGTAAAGTTAGGCCAGGGATGC
TTTGGGGAGGTGTGGATGGGCACGTGGAACGGTACTACCAGGGTTGCAATTAAGACTCTGAAAC
CCGGAACCATGTCTCCTGAGGCGTTCCTGCAAGAAGCACAAGTCATGAAGAAGCTACGTCATGA
GAAGCTAGTGCAATTGTATGCAGTTGTGAGTGAAGAGCCGATCTACATTGTCACTGAGTACATGAG
CAAGGGTTCTTTGCTGGACTTCTTGAAGGGTGAAACCGGCAAATACCTGAGACTTCCCCAGTTGG
TAGACATGGCCGCCCAGATTGCATCCGGTATGGCTTACGTGGAGAGAATGAATTACGTACACCGT
GATCTAAGAGCTGCGAACATACTGGTTGGAGAAAACTTGGTATGTAAGGTCGCTGATTTCGGTCTG
GCGAGGCTTATTGAAGACAATGAATACACTGCACGTCAAGGTGCAAAGTTCCCCATCAAGTGGAC
GGCTCCAGAGGCTGCCTTATATGGAAGGTTCACGATAAAGTCCGATGTGTGGAGTTTCGGGATAT
TGTTAACAGAATTGACAACGAAAGGACGTGTACCATATCCTGGCATGGTTAATAGAGAAGTACTTG
ACCAGGTAGAACGTGGTTATAGAATGCCATGCCCTCCGGAGTGTCCCGAGAGCTTGCACGACCTT
ATGTGTCAGTGTTGGAGGAAAGAGCCTGAGGAGAGGCCTACATTCGAGTATCTACAAGCATTCTT
AGAAGACTACTTCACGTCCACAGAACCACAGTACCAACCCGGAGAGAACTTGA

<u>Full length Src:</u>
ATGGGCAGCAATAAGTCAAAGCCGAAGGATGCAAGCCAAAGGCGTAGGTCTTTGGAGCCTGCCG
AGAATGTACATGGAGCTGGTGGTGGAGCTTTTCCGGCCAGCCAGACGCCCTCCAAACCCGCGTC
TGCTGATGGTCACCGTGGGCCAAGTGCTGCTTTTGCGCCCGCTGCAGCGGAGCCTAAGCTATTC
GGGGGTTTTAACAGTAGCGACACCGTAACGAGCCCGCAGAGAGCAGGTCCGTTGGCAGGGGGC
GTGACTACGTTCGTCGCCCTATACGACTACGAGTCTAGGACAGAGACTGACTTGAGCTTCAAAAA
GGGAGAACGTCTGCAGATCGTAAACAATACAGAGGGTGACTGGTGGCTTGCACATTCTCTTAGTA
CTGGGCAGACAGGTTATATTCCGAGCAACTATGTCGCACCGAGTGATTCAATACAGGCAGAAGAG
TGGTATTTTGGAAAAATTACTCGTAGGGAGTCCGAGAGATTATTGCTTAACGCAGAGAACCCTCGT

GGGACGTTTCTGGTCAGGGAAAGCGAAACAACAAAAGGAGCGTACTGCTTAAGCGTAAGCGATT
TCGACAATGCCAAAGGTCTTAACGTTAAGCATTATAAGATTAGGAAGTTGGACTCCGGGGGCTTTT
ATATAACGAGCAGAACCCAATTTAACTCTCTACAGCAATTGGTTGCATATTACTCCAAACATGCAGA
CGGTCTATGTCATCGTTTGACAACTGTTTGCCCCACAAGTAAGCCTCAGACGCAAGGTTTAGCAA
AGGATGCTTGGGAGATCCCTCGTGAATCACTGCGTCTTGAGGTAAAGTTAGGCCAGGGATGCTTT
GGGGAGGTGTGGATGGGCACGTGGAACGGTACTACCAGGGTTGCAATTAAGACTCTGAAACCCG
GAACCATGTCTCCTGAGGCGTTCCTGCAAGAAGCACAAGTCATGAAGAAGCTACGTCATGAGAAG
CTAGTGCAATTGTATGCAGTTGTGAGTGAAGAGCCGATCTACATTGTCACTGAGTACATGAGCAAG
GGTTCTTTGCTGGACTTCTTGAAGGGTGAAACCGGCAAATACCTGAGACTTCCCCAGTTGGTAGA
CATGGCCGCCCAGATTGCATCCGGTATGGCTTACGTGGAGAGAATGAATTACGTACACCGTGATC
TAAGAGCTGCGAACATACTGGTTGGAGAAAACTTGGTATGTAAGGTCGCTGATTTCGGTCTGGCG
AGGCTTATTGAAGACAATGAATACACTGCACGTCAAGGTGCAAAGTTCCCCATCAAGTGGACGGC
TCCAGAGGCTGCCTTATATGGAAGGTTCACGATAAAGTCCGATGTGTGGAGTTTCGGGATATTGTT
AACAGAATTGACAACGAAAGGACGTGTACCATATCCTGGCATGGTTAATAGAGAAGTACTTGACCA
GGTAGAACGTGGTTATAGAATGCCATGCCCTCCGGAGTGTCCCGAGAGCTTGCACGACCTTATGT
GTCAGTGTTGGAGGAAAGAGCCTGAGGAGAGGCCTACATTCGAGTATCTACAAGCATTCTTAGAA
GACTACTTCACGTCCACAGAACCACAGTACCAACCCGGAGAGAACTTGA

**Supplementary Table 7:** Library design

| Block | start aa | end aa | Backgrounds (neutral, gain of function, loss of function) |
|---|---|---|---|
| 1 | 268 | 326 | WT, E273G, G282A, E283L, G287P, K298M, T304A, E313R, V316I, E323R |
| 2 | 321 | 381 | WT, T341I, L328I, K354R, D368V, A370L, K354P, S345P, G355V, M377F |
| 3 | 376 | 435 | WT, E381K, R382P, R391A, I395C, T420V, R412F, Y419A, P434H, I429A |
| 4 | 431 | 491 | WT, W431V, L454A, E473T, E489S, T443W, L458V, T443P, I444F, I444P |
| 5 | 486 | 536 | WT, D496Y, E508G, Y514R, P532W, Y530D, M498Q, F523K, Y514N, L494P |

Library block 1
5' constant region:
GATGCTTGGGAGATCCCTCGT
Variable region:
GAATCACTGCGTCTTGAGGTAAAGTTAGGCCAGGGATGCTTTGGGGAGGTGTGGATGGGCACGT
GGAACGGTACTACCAGGGTTGCAATTAAGACTCTGAAACCCGGAACCATGTCTCCTGAGGCGTTC
CTGCAAGAAGCACAAGTCATGAAGAAGCTACGTCATGAGAAGCTAGTG

3' constant region:
CAATTGTATGCAGTTGTGAGTG

Library block 2
5’ constant region:
AGCACAAGTCATGAAGAAGCTA
Variable region:
CGTCATGAGAAGCTAGTGCAATTGTATGCAGTTGTGAGTGAAGAGCCGATCTACATTGTCACTGAG
TACATGAGCAAGGGTTCTTTGCTGGACTTCTTGAAGGGTGAAACCGGCAAATACCTGAGACTTCC
CCAGTTGGTAGACATGGCCGCCCAGATTGCATCCGGTATGGCTTACGTGGAG
3’ constant region:
AGAATGAATTACGTACACCGTG

Library block 3
5’ constant region:
CATGGCCGCCCAGATTGCATCC
Variable region:
GGTATGGCTTACGTGGAGAGAATGAATTACGTACACCGTGATCTAAGAGCTGCGAACATACTGGTT
GGAGAAAACTTGGTATGTAAGGTCGCTGATTTCGGTCTGGCGAGGCTTATTGAAGACAATGAATAC
ACTGCACGTCAAGGTGCAAAGTTCCCCATCAAGTGGACGGCTCCAGAG
3’ constant region:
GCTGCCTTATATGGAAGGTTCACG

Library block 4
5’ constant region:
GTGCAAAGTTCCCCATCAAG
Variable region:
TGGACGGCTCCAGAGGCTGCCTTATATGGAAGGTTCACGATAAAGTCCGATGTGTGGAGTTTCGG
GATATTGTTAACAGAATTGACAACGAAAGGACGTGTACCATATCCTGGCATGGTTAATAGAGAAGTA
CTTGACCAGGTAGAACGTGGTTATAGAATGCCATGCCCTCCGGAGTGTCCC

3’ constant region:
GAGAGCTTGCACGACCTTATG

Library block 5
5’ constant region:
GAACGTGGTTATAGAATGCCA
Variable region:
TGCCCTCCGGAGTGTCCCGAGAGCTTGCACGACCTTATGTGTCAGTGTTGGAGGAAAGAGCCTG
AGGAGAGGCCTACATTCGAGTATCTACAAGCATTCTTAGAAGACTACTTCACGTCCACAGAACCAC
AGTACCAACCCGGAGAGAACTTG
3’ constant region:
AAGCTTTAAATTAGTTATGTCACGC

Genetic background sequences:

Block 1:
>WT
GATGCTTGGGAGATCCCTCGTGAATCACTGCGTCTTGAGGTAAAGTTAGGCCAGGGATGCTTTGG
GGAGGTGTGGATGGGCACGTGGAACGGTACTACCAGGGTTGCAATTAAGACTCTGAAACCCGGA
ACCATGTCTCCTGAGGCGTTCCTGCAAGAAGCACAAGTCATGAAGAAGCTACGTCATGAGAAGCT
AGTGCAATTGTATGCAGTTGTGAGTG
>E273G

GATGCTTGGGAGATCCCTCGTGAATCACTGCGTCTTGGTGTAAAGTTAGGCCAGGGATGCTTTGG
GGAGGTGTGGATGGGCACGTGGAACGGTACTACCAGGGTTGCAATTAAGACTCTGAAACCCGGA
ACCATGTCTCCTGAGGCGTTCCTGCAAGAAGCACAAGTCATGAAGAAGCTACGTCATGAGAAGCT
AGTGCAATTGTATGCAGTTGTGAGTG
>G282A
GATGCTTGGGAGATCCCTCGTGAATCACTGCGTCTTGAGGTAAAGTTAGGCCAGGGATGCTTTGC
AGAGGTGTGGATGGGCACGTGGAACGGTACTACCAGGGTTGCAATTAAGACTCTGAAACCCGGA
ACCATGTCTCCTGAGGCGTTCCTGCAAGAAGCACAAGTCATGAAGAAGCTACGTCATGAGAAGCT
AGTGCAATTGTATGCAGTTGTGAGTG
>E283L
GATGCTTGGGAGATCCCTCGTGAATCACTGCGTCTTGAGGTAAAGTTAGGCCAGGGATGCTTTGG
GTTAGTGTGGATGGGCACGTGGAACGGTACTACCAGGGTTGCAATTAAGACTCTGAAACCCGGAA
CCATGTCTCCTGAGGCGTTCCTGCAAGAAGCACAAGTCATGAAGAAGCTACGTCATGAGAAGCTA
GTGCAATTGTATGCAGTTGTGAGTG
>G287P
GATGCTTGGGAGATCCCTCGTGAATCACTGCGTCTTGAGGTAAAGTTAGGCCAGGGATGCTTTGG
GGAGGTGTGGATGCCAACGTGGAACGGTACTACCAGGGTTGCAATTAAGACTCTGAAACCCGGA
ACCATGTCTCCTGAGGCGTTCCTGCAAGAAGCACAAGTCATGAAGAAGCTACGTCATGAGAAGCT
AGTGCAATTGTATGCAGTTGTGAGTG
>K298M
GATGCTTGGGAGATCCCTCGTGAATCACTGCGTCTTGAGGTAAAGTTAGGCCAGGGATGCTTTGG
GGAGGTGTGGATGGGCACGTGGAACGGTACTACCAGGGTTGCAATTATGACTCTGAAACCCGGA
ACCATGTCTCCTGAGGCGTTCCTGCAAGAAGCACAAGTCATGAAGAAGCTACGTCATGAGAAGCT
AGTGCAATTGTATGCAGTTGTGAGTG
>T304A
GATGCTTGGGAGATCCCTCGTGAATCACTGCGTCTTGAGGTAAAGTTAGGCCAGGGATGCTTTGG
GGAGGTGTGGATGGGCACGTGGAACGGTACTACCAGGGTTGCAATTAAGACTCTGAAACCCGGA
GCAATGTCTCCTGAGGCGTTCCTGCAAGAAGCACAAGTCATGAAGAAGCTACGTCATGAGAAGCT
AGTGCAATTGTATGCAGTTGTGAGTG
>E313R
GATGCTTGGGAGATCCCTCGTGAATCACTGCGTCTTGAGGTAAAGTTAGGCCAGGGATGCTTTGG
GGAGGTGTGGATGGGCACGTGGAACGGTACTACCAGGGTTGCAATTAAGACTCTGAAACCCGGA
ACCATGTCTCCTGAGGCGTTCCTGCAAAGAGCACAAGTCATGAAGAAGCTACGTCATGAGAAGCT
AGTGCAATTGTATGCAGTTGTGAGTG
>V316I
GATGCTTGGGAGATCCCTCGTGAATCACTGCGTCTTGAGGTAAAGTTAGGCCAGGGATGCTTTGG
GGAGGTGTGGATGGGCACGTGGAACGGTACTACCAGGGTTGCAATTAAGACTCTGAAACCCGGA
ACCATGTCTCCTGAGGCGTTCCTGCAAGAAGCACAAATTATGAAGAAGCTACGTCATGAGAAGCT
AGTGCAATTGTATGCAGTTGTGAGTG
>E323R
GATGCTTGGGAGATCCCTCGTGAATCACTGCGTCTTGAGGTAAAGTTAGGCCAGGGATGCTTTGG
GGAGGTGTGGATGGGCACGTGGAACGGTACTACCAGGGTTGCAATTAAGACTCTGAAACCCGGA
ACCATGTCTCCTGAGGCGTTCCTGCAAGAAGCACAAGTCATGAAGAAGCTACGTCATAGAAAGCT
AGTGCAATTGTATGCAGTTGTGAGTG

Block 2:
>WT
CGTCATGAGAAGCTAGTGCAATTGTATGCAGTTGTGAGTGAAGAGCCGATCTACATTGTCACTGAG
TACATGAGCAAGGGTTCTTTGCTGGACTTCTTGAAGGGTGAAACCGGCAAATACCTGAGACTTCC
CCAGTTGGTAGACATGGCCGCCCAGATTGCATCCGGTATGGCTTACGTGGAG
>L328I

CGTCATGAGAAGCTAGTGCAAATATATGCAGTTGTGAGTGAAGAGCCGATCTACATTGTCACTGAG
TACATGAGCAAGGGTTCTTTGCTGGACTTCTTGAAGGGTGAAACCGGCAAATACCTGAGACTTCC
CCAGTTGGTAGACATGGCCGCCCAGATTGCATCCGGTATGGCTTACGTGGAG
>T341I
CGTCATGAGAAGCTAGTGCAATTGTATGCAGTTGTGAGTGAAGAGCCGATCTACATTGTCATAGAG
TACATGAGCAAGGGTTCTTTGCTGGACTTCTTGAAGGGTGAAACCGGCAAATACCTGAGACTTCC
CCAGTTGGTAGACATGGCCGCCCAGATTGCATCCGGTATGGCTTACGTGGAG
>K354R
CGTCATGAGAAGCTAGTGCAATTGTATGCAGTTGTGAGTGAAGAGCCGATCTACATTGTCACTGAG
TACATGAGCAAGGGTTCTTTGCTGGACTTCTTGAGAGGTGAAACCGGCAAATACCTGAGACTTCC
CCAGTTGGTAGACATGGCCGCCCAGATTGCATCCGGTATGGCTTACGTGGAG
>D368V
CGTCATGAGAAGCTAGTGCAATTGTATGCAGTTGTGAGTGAAGAGCCGATCTACATTGTCACTGAG
TACATGAGCAAGGGTTCTTTGCTGGACTTCTTGAAGGGTGAAACCGGCAAATACCTGAGACTTCC
CCAGTTGGTAGTGATGGCCGCCCAGATTGCATCCGGTATGGCTTACGTGGAG
>A370L
CGTCATGAGAAGCTAGTGCAATTGTATGCAGTTGTGAGTGAAGAGCCGATCTACATTGTCACTGAG
TACATGAGCAAGGGTTCTTTGCTGGACTTCTTGAAGGGTGAAACCGGCAAATACCTGAGACTTCC
CCAGTTGGTAGACATGTTGGCCCAGATTGCATCCGGTATGGCTTACGTGGAG
>K354P
CGTCATGAGAAGCTAGTGCAATTGTATGCAGTTGTGAGTGAAGAGCCGATCTACATTGTCACTGAG
TACATGAGCAAGGGTTCTTTGCTGGACTTCTTGCCCGGTGAAACCGGCAAATACCTGAGACTTCC
CCAGTTGGTAGACATGGCCGCCCAGATTGCATCCGGTATGGCTTACGTGGAG
>S345P
CGTCATGAGAAGCTAGTGCAATTGTATGCAGTTGTGAGTGAAGAGCCGATCTACATTGTCACTGAG
TACATGCCAAAGGGTTCTTTGCTGGACTTCTTGAAGGGTGAAACCGGCAAATACCTGAGACTTCC
CCAGTTGGTAGACATGGCCGCCCAGATTGCATCCGGTATGGCTTACGTGGAG
>G355V
CGTCATGAGAAGCTAGTGCAATTGTATGCAGTTGTGAGTGAAGAGCCGATCTACATTGTCACTGAG
TACATGAGCAAGGGTTCTTTGCTGGACTTCTTGAAGGTCGAAACCGGCAAATACCTGAGACTTCC
CCAGTTGGTAGACATGGCCGCCCAGATTGCATCCGGTATGGCTTACGTGGAG
>M377F
CGTCATGAGAAGCTAGTGCAATTGTATGCAGTTGTGAGTGAAGAGCCGATCTACATTGTCACTGAG
TACATGAGCAAGGGTTCTTTGCTGGACTTCTTGAAGGGTGAAACCGGCAAATACCTGAGACTTCC
CCAGTTGGTAGACATGGCCGCCCAGATTGCATCCGGTTTTGCTTACGTGGAG

Block 3:
>WT
GGTATGGCTTACGTGGAGAGAATGAATTACGTACACCGTGATCTAAGAGCTGCGAACATACTGGTT
GGAGAAAACTTGGTATGTAAGGTCGCTGATTTCGGTCTGGCGAGGCTTATTGAAGACAATGAATAC
ACTGCACGTCAAGGTGCAAAGTTCCCCATCAAGTGGACGGCTCCAGAG
>E381K
GGTATGGCTTACGTGAAAAGAATGAATTACGTACACCGTGATCTAAGAGCTGCGAACATACTGGTT
GGAGAAAACTTGGTATGTAAGGTCGCTGATTTCGGTCTGGCGAGGCTTATTGAAGACAATGAATAC
ACTGCACGTCAAGGTGCAAAGTTCCCCATCAAGTGGACGGCTCCAGAG
>R382P
GGTATGGCTTACGTGGAGCCCATGAATTACGTACACCGTGATCTAAGAGCTGCGAACATACTGGTT
GGAGAAAACTTGGTATGTAAGGTCGCTGATTTCGGTCTGGCGAGGCTTATTGAAGACAATGAATAC
ACTGCACGTCAAGGTGCAAAGTTCCCCATCAAGTGGACGGCTCCAGAG
>R391A

GGTATGGCTTACGTGGAGAGAATGAATTACGTACACCGTGATCTAGCCGCTGCGAACATACTGGTT
GGAGAAAACTTGGTATGTAAGGTCGCTGATTTCGGTCTGGCGAGGCTTATTGAAGACAATGAATAC
ACTGCACGTCAAGGTGCAAAGTTCCCCATCAAGTGGACGGCTCCAGAG
>I395C
GGTATGGCTTACGTGGAGAGAATGAATTACGTACACCGTGATCTAAGAGCTGCGAACTGCCTGGT
TGGAGAAAACTTGGTATGTAAGGTCGCTGATTTCGGTCTGGCGAGGCTTATTGAAGACAATGAATA
CACTGCACGTCAAGGTGCAAAGTTCCCCATCAAGTGGACGGCTCCAGAG
>T420V
GGTATGGCTTACGTGGAGAGAATGAATTACGTACACCGTGATCTAAGAGCTGCGAACATACTGGTT
GGAGAAAACTTGGTATGTAAGGTCGCTGATTTCGGTCTGGCGAGGCTTATTGAAGACAATGAATAC
GTCGCACGTCAAGGTGCAAAGTTCCCCATCAAGTGGACGGCTCCAGAG
>R412F
GGTATGGCTTACGTGGAGAGAATGAATTACGTACACCGTGATCTAAGAGCTGCGAACATACTGGTT
GGAGAAAACTTGGTATGTAAGGTCGCTGATTTCGGTCTGGCGTTTCTTATTGAAGACAATGAATAC
ACTGCACGTCAAGGTGCAAAGTTCCCCATCAAGTGGACGGCTCCAGAG
>Y419A
GGTATGGCTTACGTGGAGAGAATGAATTACGTACACCGTGATCTAAGAGCTGCGAACATACTGGTT
GGAGAAAACTTGGTATGTAAGGTCGCTGATTTCGGTCTGGCGAGGCTTATTGAAGACAATGAAGC
AACTGCACGTCAAGGTGCAAAGTTCCCCATCAAGTGGACGGCTCCAGAG
>P434H
GGTATGGCTTACGTGGAGAGAATGAATTACGTACACCGTGATCTAAGAGCTGCGAACATACTGGTT
GGAGAAAACTTGGTATGTAAGGTCGCTGATTTCGGTCTGGCGAGGCTTATTGAAGACAATGAATAC
ACTGCACGTCAAGGTGCAAAGTTCCCCATCAAGTGGACGGCTCACGAG
>I429A
GGTATGGCTTACGTGGAGAGAATGAATTACGTACACCGTGATCTAAGAGCTGCGAACATACTGGTT
GGAGAAAACTTGGTATGTAAGGTCGCTGATTTCGGTCTGGCGAGGCTTATTGAAGACAATGAATAC
ACTGCACGTCAAGGTGCAAAGTTCCCCGCGAAGTGGACGGCTCCAGAG

Block 4:
>WT
TGGACGGCTCCAGAGGCTGCCTTATATGGAAGGTTCACGATAAAGTCCGATGTGTGGAGTTTCGG
GATATTGTTAACAGAATTGACAACGAAAGGACGTGTACCATATCCTGGCATGGTTAATAGAGAAGTA
CTTGACCAGGTAGAACGTGGTTATAGAATGCCATGCCCTCCGGAGTGTCCC
>W431V
GTCACGGCTCCAGAGGCTGCCTTATATGGAAGGTTCACGATAAAGTCCGATGTGTGGAGTTTCGG
GATATTGTTAACAGAATTGACAACGAAAGGACGTGTACCATATCCTGGCATGGTTAATAGAGAAGTA
CTTGACCAGGTAGAACGTGGTTATAGAATGCCATGCCCTCCGGAGTGTCCC
>L454A
TGGACGGCTCCAGAGGCTGCCTTATATGGAAGGTTCACGATAAAGTCCGATGTGTGGAGTTTCGG
GATAGCTTTAACAGAATTGACAACGAAAGGACGTGTACCATATCCTGGCATGGTTAATAGAGAAGT
ACTTGACCAGGTAGAACGTGGTTATAGAATGCCATGCCCTCCGGAGTGTCCC
>E473T
TGGACGGCTCCAGAGGCTGCCTTATATGGAAGGTTCACGATAAAGTCCGATGTGTGGAGTTTCGG
GATATTGTTAACAGAATTGACAACGAAAGGACGTGTACCATATCCTGGCATGGTTAATAGAACTGTA
CTTGACCAGGTAGAACGTGGTTATAGAATGCCATGCCCTCCGGAGTGTCCC
>E489S
TGGACGGCTCCAGAGGCTGCCTTATATGGAAGGTTCACGATAAAGTCCGATGTGTGGAGTTTCGG
GATATTGTTAACAGAATTGACAACGAAAGGACGTGTACCATATCCTGGCATGGTTAATAGAGAAGTA
CTTGACCAGGTAGAACGTGGTTATAGAATGCCATGCCCTCCGTCTTGTCCC
>T443W

TGGACGGCTCCAGAGGCTGCCTTATATGGAAGGTTCTGGATAAAGTCCGATGTGTGGAGTTTCGG
GATATTGTTAACAGAATTGACAACGAAAGGACGTGTACCATATCCTGGCATGGTTAATAGAGAAGTA
CTTGACCAGGTAGAACGTGGTTATAGAATGCCATGCCCTCCGGAGTGTCCC
>L458V
TGGACGGCTCCAGAGGCTGCCTTATATGGAAGGTTCACGATAAAGTCCGATGTGTGGAGTTTCGG
GATATTGTTAACAGAAGTAACAACGAAAGGACGTGTACCATATCCTGGCATGGTTAATAGAGAAGTA
CTTGACCAGGTAGAACGTGGTTATAGAATGCCATGCCCTCCGGAGTGTCCC
>T443P
TGGACGGCTCCAGAGGCTGCCTTATATGGAAGGTTCCCAATAAAGTCCGATGTGTGGAGTTTCGG
GATATTGTTAACAGAATTGACAACGAAAGGACGTGTACCATATCCTGGCATGGTTAATAGAGAAGTA
CTTGACCAGGTAGAACGTGGTTATAGAATGCCATGCCCTCCGGAGTGTCCC
>I444F
TGGACGGCTCCAGAGGCTGCCTTATATGGAAGGTTCACGTTCAAGTCCGATGTGTGGAGTTTCGG
GATATTGTTAACAGAATTGACAACGAAAGGACGTGTACCATATCCTGGCATGGTTAATAGAGAAGTA
CTTGACCAGGTAGAACGTGGTTATAGAATGCCATGCCCTCCGGAGTGTCCC
>I444P
TGGACGGCTCCAGAGGCTGCCTTATATGGAAGGTTCACGCCGAAGTCCGATGTGTGGAGTTTCG
GGATATTGTTAACAGAATTGACAACGAAAGGACGTGTACCATATCCTGGCATGGTTAATAGAGAAG
TACTTGACCAGGTAGAACGTGGTTATAGAATGCCATGCCCTCCGGAGTGTCCC

Block 5:
>WT
TGCCCTCCGGAGTGTCCCGAGAGCTTGCACGACCTTATGTGTCAGTGTTGGAGGAAAGAGCCTG
AGGAGAGGCCTACATTCGAGTATCTACAAGCATTCTTAGAAGACTACTTCACGTCCACAGAACCAC
AGTACCAACCCGGAGAGAACTTG
>D496Y
TGCCCTCCGGAGTGTCCCGAGAGCTTGCACTATCTTATGTGTCAGTGTTGGAGGAAAGAGCCTG
AGGAGAGGCCTACATTCGAGTATCTACAAGCATTCTTAGAAGACTACTTCACGTCCACAGAACCAC
AGTACCAACCCGGAGAGAACTTG
>E508G
TGCCCTCCGGAGTGTCCCGAGAGCTTGCACGACCTTATGTGTCAGTGTTGGAGGAAAGAGCCTG
AGGGCAGGCCTACATTCGAGTATCTACAAGCATTCTTAGAAGACTACTTCACGTCCACAGAACCAC
AGTACCAACCCGGAGAGAACTTG
>Y514R
TGCCCTCCGGAGTGTCCCGAGAGCTTGCACGACCTTATGTGTCAGTGTTGGAGGAAAGAGCCTG
AGGAGAGGCCTACATTCGAGAGGCTACAAGCATTCTTAGAAGACTACTTCACGTCCACAGAACCA
CAGTACCAACCCGGAGAGAACTTG
>P532W
TGCCCTCCGGAGTGTCCCGAGAGCTTGCACGACCTTATGTGTCAGTGTTGGAGGAAAGAGCCTG
AGGAGAGGCCTACATTCGAGTATCTACAAGCATTCTTAGAAGACTACTTCACGTCCACAGAACCAC
AGTACCAATGGGGAGAGAACTTG
>Y530D
TGCCCTCCGGAGTGTCCCGAGAGCTTGCACGACCTTATGTGTCAGTGTTGGAGGAAAGAGCCTG
AGGAGAGGCCTACATTCGAGTATCTACAAGCATTCTTAGAAGACTACTTCACGTCCACAGAACCAC
AGGATCAACCCGGAGAGAACTTG
>M498Q
TGCCCTCCGGAGTGTCCCGAGAGCTTGCACGACCTTCAATGTCAGTGTTGGAGGAAAGAGCCTG
AGGAGAGGCCTACATTCGAGTATCTACAAGCATTCTTAGAAGACTACTTCACGTCCACAGAACCAC
AGTACCAACCCGGAGAGAACTTG
>F523K

TGCCCTCCGGAGTGTCCCGAGAGCTTGCACGACCTTATGTGTCAGTGTTGGAGGAAAGAGCCTG
AGGAGAGGCCTACATTCGAGTATCTACAAGCATTCTTAGAAGACTACAAAACGTCCACAGAACCAC
AGTACCAACCCGGAGAGAACTTG
>Y514N
TGCCCTCCGGAGTGTCCCGAGAGCTTGCACGACCTTATGTGTCAGTGTTGGAGGAAAGAGCCTG
AGGAGAGGCCTACATTCGAGAACCTACAAGCATTCTTAGAAGACTACTTCACGTCCACAGAACCA
CAGTACCAACCCGGAGAGAACTTG
>L494P
TGCCCTCCGGAGTGTCCCGAGAGCCCACACGACCTTATGTGTCAGTGTTGGAGGAAAGAGCCTG
AGGAGAGGCCTACATTCGAGTATCTACAAGCATTCTTAGAAGACTACTTCACGTCCACAGAACCAC
AGTACCAACCCGGAGAGAACTTG

**Supplementary Table 8:** Kinase Atlas known allosteric pockets from the literature

| Site | Site Name Origin | Inhibitor Type | Source Kinase | PDB | Pocket Description | Present in Src |
|---|---|---|---|---|---|---|
| DFG | DFG motif | II | many | 1IEP | Hydrophobic pocket that opens up when DFG motif switches to inactive "DFG-out" conformation; binding here may stabilize inactive kinase conformation | yes |
| MT3 | MEK1/2 type III inhibitor | III | MEK1/2 | 4AN2 | Adjacent to ATP and DFG-out pockets; binding disrupts salt bridge required for kinase activity | yes |
| PIF | PDK1 interacting fragment | IV | PDK1 | 4RQK | PDK1 regulates other AGC kinases by recruiting them through this site | no |
| MPP | MKK4 p38a peptide | IV | MKK4 | 3ALO | p38a peptide binding inhibits MKK4 by inducing conformational changes that lead to auto-inhibition | yes |
| CMP | c-Abl myristoyl pocket | IV | c-Abl | 3K5V | Binding here leads to active or inactive state in c-Abl (depending on ligand size) by affecting SH domain binding | yes |
| PMP | PKA myristoyl pocket | IV | PKA | 1CMK | Myristoyl binding here activates membrane binding in PKA | no |
| DRS | D-recruitment site | IV | all MAPKs | 1UKI | Substrate docking site present in all MAP kinases | no |
| DEF | docking site for ERK FXF | IV | some MAPKs | 3O2M | Substrate docking site present in some MAP kinases; located near MAPK insert | no |

| | | | | | | |
|---|---|---|---|---|---|---|
| LBP | lipid binding pocket | IV | p38a MAPK | 3NEW | Binding of different lipids here affects p38a MAPK's preference and activity for different substrates | no |
| PDIG | PDIG motif | IV | Chk1 | 3JVS | Substrate recognition site located near PDIG motif in Chk1 | yes |
| AAS | Aurora A activation segment | IV | Aurora A | 4C3P | An Aurora A monomer activates another through binding of its activation segment to this site | yes |
| EDI | EGFR dimerization interface | IV | EGFR | 2RFE | An EGFR monomer activates another by binding at this interface on the C-terminal domain | yes |
